# Advanced Age in Mice Exacerbates Sepsis-Induced Inflammation, Vascular Permeability, and Multi-Organ Dysfunction

**DOI:** 10.1101/2025.05.27.656343

**Authors:** Han Noo Ri Lee, Jason Lin, Camryn J. Smith, Lorraine B. Ware, Fiona E. Harrison, Julie A. Bastarache, Brandon Baer

## Abstract

Sepsis is a life-threatening syndrome marked by a dysregulated immune response to an infection and significant endothelial vascular permeability, often leading to multi-organ failure. Elderly patients are particularly vulnerable to sepsis, with higher morbidity and mortality rates. We hypothesized that advanced age exacerbates sepsis-induced inflammation and endothelial vascular permeability, resulting in a delayed recovery, persistent inflammation, and sustained organ injury. Using a polymicrobial sepsis model in young (3-month-old) and aged (18-month-old) C57BL/6 mice, sepsis was induced via intraperitoneal cecal slurry (CS) injection. Outcomes were assessed during the acute (24-hour; 1.6mg/g CS) and recovery (8-day; 1.0 mg/g CS) phases. During the acute phase, aged mice exhibited worse physiologic dysfunction, higher systemic (plasma TNF-a: young septic 202.1 pg/mL [17.44, 398.9] vs. aged septic 482.6 pg/mL [279.8, 711.7]; p = 0.0352 Mann-Whitney) and organ-specific inflammation, increased endothelial injury and vascular permeability, as well as greater kidney and liver dysfunction compared to young mice. During recovery, aged mice had sustained physiologic dysfunction, prolonged systemic and organ-specific inflammation, and sustained organ injury (kidney tissue NGAL: young septic 291.5 RE [203.7, 373.2] vs. aged septic 821 RE [456, 1258] protein normalized to beta actin; p = 0.0008 Mann-Whitney) compared to young mice. These results support the hypothesis that advanced age worsens sepsis severity and outcomes and delays recovery, emphasizing the need for aged models and multi-organ evaluations to develop effective therapies for this vulnerable population.

## INTRODUCTION

Sepsis is an increasingly common and serious syndrome, particularly among the elderly, who face significantly higher rates of morbidity and mortality, with no specific therapy or means of prevention ^1,2^. Characterized by a dysregulated host response to infection and endothelial vascular barrier permeability, sepsis in the elderly often leads to life-threatening organ dysfunction in the lungs, kidneys, and liver ^3–9^. Adults over 65 currently represent 65% of all septic patients, 67% of septic patients with organ failure, and 75% of sepsis-related deaths ^10–18^. However, most animal studies modeling sepsis utilize young adult mice (8 to 12 weeks old), corresponding to adolescent and early adulthood in humans – an age group with the lowest risk of developing sepsis and the highest likelihood of recovering without issue ^19–22^. Thus, there is a clear need for research using aged animals to better our understanding of sepsis pathogenesis in this large and high-risk demographic.

Sepsis is a complex syndrome involving multiple organ systems, whose development, progression, and severity can vary widely among individual patients ^2,13,23–26^. This ranges from patients in which sepsis leads to multiple organ dysfunction and ultimately death, to patients who recover quickly with few lasting effects. Persistent inflammation and vascular permeability are at the core of the more severe sepsis pathophysiology ^1,27–31^. While the activation of pro-inflammatory processes is essential to combat infection, under septic conditions, persistence of this pro-inflammatory state becomes harmful ^1,3,32,33^. This maladaptive immune response is often linked to increased endothelial damage and barrier permeability that cause functional impairments in multiple organs ^28,29,34^. Notably, inflammation and barrier permeability-induced impairments in the lungs, liver, and kidneys are significant contributors to sepsis mortality and morbidity ^25,26,34,35^.

Aging independently exacerbates many of the pathological processes associated with severe sepsis ^10,11,14–17,36^. Advanced age (>65 years) is linked to heightened baseline inflammation, an excessive release of cytokines in response to acute sepsis, and worse overall recovery from sepsis ^15,37–40^. Additionally, advanced age contributes to vascular endothelial cell dysfunction and increased vascular permeability in the lungs, kidneys, and liver, leading to greater organ dysfunction ^41–44^. However, few studies have examined the impact of advanced age on recovery from sepsis. While the effects of acute sepsis and aging on inflammation, vascular endothelial cell barrier dysfunction, and organ dysfunction have been previously investigated separately ^45–53^, there is a need for a better understanding of how these factors interact during the recovery phase of sepsis and how they influence mortality and illness severity. We hypothesized that advanced age would worsen sepsis-induced inflammation and endothelial vascular barrier permeability, leading to delayed recovery, persistent inflammation, and sustained organ injury.

## METHODS

### Animals

All animal studies were approved by the Vanderbilt University Medical Center (VUMC) Institutional Animal Care and Use Committee (Protocol number: M2200022-00). Male and female wildtype C57BL/6 mice, young (3-month-old) and aged (18-month-old), were obtained from the National Institute on Aging (NIA). Mice were allowed to acclimatize in the VUMC animal facility for at least one week and were housed in a temperature-controlled room (12:12-hour light/dark cycles) with free access to water and standard food.

### Polymicrobial Sepsis Model of Cecal Slurry

Polymicrobial peritoneal sepsis was induced using cecal slurry (CS), a model that has been described previously ^54–59^. Briefly, CS was prepared by mixing cecal contents collected from female C57BL/6 donor mice (6 weeks of age) purchased from The Jackson Laboratory (Bar Harbor, ME). Mice were euthanized within 7 days of arrival at VUMC, with their cecal contents collected, resuspended in 5% dextrose (80 mg/mL), filtered through a 70-mm cell strainer, and resuspended in glycerol. To minimize variability, a large batch of CS was prepared, with aliquots stored at −80°C.

To induce sepsis, recipient mice were intraperitoneally injected with CS. To ensure an adequate sample size and observable sepsis-induced illness at both the acute (24-hours) and recovery (8-days) phases of sepsis, the survival rate and sepsis illness scores were measured in a separate cohort of young (N=25) and aged (N=44) mice injected with different doses of CS (1.0-2.0 mg/g of body weight). Based on the findings from this initial cohort, mice were given 1.6 mg/g CS by body weight for all subsequent acute phase sepsis experiments with endpoints up to 24 hours after infection, while mice were given 1.0 mg/g CS for all recovery phase sepsis experiments with endpoints up to 8-days after infection. Mice were administered 300 µL of intraperitoneal antibiotics (imipenem and cilastatin NDC 63323-349-01; 50 mg/mL) and 700 µL of subcutaneous sterile saline 12 hours after CS injection ^59^ in the acute phase experiments, while those in the recovery phase experiments received the same injections at both 6 and 12 hours. Control mice were administered 5% dextrose intraperitoneally dose-matched to their experimental counterparts as well as antibiotics and fluids.

### Tissue Collection

Mice were euthanized with intraperitoneal injection of sodium pentobarbital (Sigma-Aldrich, St. Louis, MO) at 24-hours or 8-days after CS injection. Blood was collected in K_2_EDTA (Becton, Dickinson and Company, Franklin Lakes, NJ) tubes by blind cardiac puncture with a 25-gauge needle. Centrifugation at 1500 x g for 10 minutes at 4°C was used to isolate plasma.

The left mainstem bronchus was ligated and a bronchoalveolar lavage (BAL) with 600 µL of sterile saline was performed on right lung. BAL was centrifuged at 1500 x g for 10 minutes at 4°C. Supernatant was collected and stored at −80°C. The left lung was removed, weighed, and dried at 55°C for 72 hours to determine wet-to-dry weight ratio.

Kidney and liver tissue were collected, flash-frozen in liquid nitrogen, and stored at −80°C. The spleen was collected, weighed, and briefly submerged in 70% ethanol to remove adherent bacteria before being transferred to a homogenization tube with 2.8 mm ceramic beads (Fisher Scientific, Waltham, MA) containing 1 mL of sterile phosphate-buffered saline (PBS). Spleens were homogenized at 4.85 m/s for 20 seconds using Bead Mill 24 homogenizer (Fisher Scientific, Waltham, MA). Spleen homogenate was then serially diluted in PBS, plated onto Lysogeny broth agar, and incubated overnight at 37°C. Colony forming units (CFU) were counted the following day.

### Experimental Outcomes

#### Illness Severity, Mortality, Body Weight, Body Temperature, and Metabolism

Illness severity was assessed in a blinded fashion using a validated semi-quantitative sepsis scoring system on mixed cages that contained both control and septic mice ^57,60,61^. Briefly, mice were scored out of 12 with points being deducted based on lack of responsiveness and physical appearance parameters of illness. A score of 12 represented a healthy mouse with normal behavior, a score of 0 represented a dead mouse, and a humane endpoint for this sepsis scoring system was set at 4 or below. Sepsis scoring was conducted every 4 hours for the acute phase experiments. For the recovery phase experiments, sepsis scores were evaluated every 3-4 hours for the first 36 hours, then every 12 hours until mice achieved a score of 12, after which they were scored every 24 hours. Body weight and temperature (Contec Infrared Thermometer model TP500; Contec Co., Ltd. Osaka, Japan) were also measured at these timepoints. Physiologic dysfunction was assessed through changes in body weight, body temperature, plasma glucose, plasma cholesterol, and plasma triglycerides. Body weights were recorded before CS injections and at the 24-hour or 8-day timepoints, respectively. For the acute phase, body temperatures were recorded every 4 hours, while for the recovery phase they were recorded every 6 hours for the first 12 hours and every 12-24 hours thereafter. Blood chemistry analysis was performed by the Translational Pathology Shared Resource Core (Nashville, TN).

#### Systemic Inflammation

To evaluate systemic inflammation, plasma concentrations of interferon gamma (IFN-γ), interleukin-10 (IL-10), interleukin-1 beta (IL-1β), interleukin-6 (IL-6), C-X-C motif chemokine ligand 1 (CXCL1), and tumor necrosis factor alpha (TNF-α) were measured using a V-PLEX Proinflammatory Panel 1 (Meso Scale Diagnostics, Rockville, MD).

#### Lung Inflammation

To evaluate lung inflammation, cell counts, differentials, and inflammatory cytokine concentrations were measured in the BAL. Cell counts and differentials were performed on fresh BAL using a light microscope and hemocytometer. BAL concentrations of IFN-γ, IL-10, IL-1β, IL-6, CXCL1, and TNF-α were measured using V-PLEX Proinflammatory Panel 1 (Meso Scale Diagnostics, Rockville, MD).

#### Alveolar-capillary Barrier Permeability

Alveolar-capillary barrier permeability was evaluated using BAL total protein concentration quantified using Pierce bicinchoninic acid assay (Thermo Scientific, Waltham, MA).

#### Liver and Kidney Inflammation

Liver and kidney inflammation were assessed through gene expression levels of pro-inflammatory cytokines in kidney or liver tissue. Briefly, gene expression was measured by qPCR using frozen tissue, a Qiagen RNeasy Plus Mini Kit (Hilden, Germany) to isolate mRNA, and an iScript cDNA Synthesis Kit (Bio-Rad, Hercules, CA) to synthesize cDNA. PrimePCR probes (Bio-rad, Hercules, CA) were used to measure IFN-γ (10031230), IL-10 (10031227), IL-1β (10031230), IL-6 (10031227), CXCL1 (10031227), and TNF-α (10031227) expression normalized to GAPDH (10031226) expression.

#### Liver and Kidney Dysfunction and Injury

Liver dysfunction was assessed using plasma concentrations of total bilirubin and albumin, while liver injury was measured through plasma concentrations of alanine aminotransferase (ALT) and gamma-glutamyl transferase (GGT). Kidney dysfunction was evaluated by plasma urea nitrogen concentrations, while kidney injury was assessed through gene and protein expression levels of kidney injury molecule-1 (KIM-1) and neutrophil gelatinase associated lipocalin (NGAL) in kidney tissue. Plasma urea nitrogen concentrations were measured using a Urea Nitrogen Colorimetric Detection Kit (Invitrogen, Waltham, MA). KIM-1 and NGAL protein levels were measured by Western blot using frozen kidney tissue homogenate prepared in RIPA buffer (Thermo Scientific, Waltham, MA). Briefly, the membrane was blocked in Intercept Blocking Buffer (Li-Cor, Lincoln, NB) before incubation in KIM-1 (1:250) antibody (Goat AF1817, R&D Systems, Minneapolis, MN) or NGAL (1:1000) antibody (Goat AF1857, R&D Systems, Minneapolis, MN) for 24 hours. Li-Cor Odyssey Imager was used to analyze the blots and target protein level was normalized to β-actin (1:20,000; Cell Signaling Technology, Danvers, MA; Rabbit 4970)n. Gene expression was measured by qPCR using frozen kidney tissue. Briefly, Qiagen RNeasy Plus Mini Kit (Hilden, Germany) was used to isolate mRNA, while iScript cDNA Synthesis Kit (Bio-Rad, Hercules, CA) was used to synthesize cDNA. NGAL (Lcn2; 10031226) and KIM-1 (Havcr1; 10031229) expression were measured using PrimePCR probes (Bio-Rad, Hercules, CA), and normalized to GAPDH (10031226) expression.

#### Endothelial Injury and Vascular Permeability

Endothelial injury and vascular permeability were assessed through Angiosense and plasma p-selectin concentrations, respectively. Briefly, mice were given a retro-orbital injection of 100 µL IVISense Vascular 750 Fluorescent Probe (AngioSense; 2 nmol per 100 µL; Revvity, Waltham, MA) at the same time as intraperitoneal CS or 5% dextrose injection. Accumulation of AngioSense in whole lung and kidney was measured using a Li-Cor Pearl camera. Plasma soluble p-selectin concentrations were measured using mouse sP-Selectin/CD62P Quantikine ELISA kit (R&D Systems, Minneapolis, MN).

#### Data Analysis

Across all studies, animals were excluded from analysis if: 1) the sepsis illness score of animals receiving CS did not drop below 12, indicating they did not become ill (N = 6 young, N = 2 aged) or 2) the sepsis illness score of animals receiving 5% dextrose dropped below 11, indicating sickness from non-experimental cause (N = 1 aged). Data were analyzed using GraphPad Prism 9.5.1. and values were also excluded for specific outcomes if identified as an outlier (Q=0.1%) via a robust regression and outlier removal (ROUT) outlier test within their own groups ^62^. Data from males (N = 26 young, 34 aged) and females (N = 24 young, 39 aged) were analyzed together and either presented as individual data points (separate symbols for male and female mice) with their combined median, combined median +/− interquartile range, or as a combined mean +/− standard error of the mean. Continuous variables that were not normally distributed were log-transformed prior to analysis with a two-way ANOVA and Tukey’s multiple comparison or uncorrected Fisher’s LSD post-hoc test. In the case where log-transformation was not feasible due to data points with a value of zero or a negative value, a Kruskal-Wallis test and Dunn’s post-hoc test were utilized. A p-value of less than 0.05 was considered statistically significant, with a Bonferroni correction being used for sex-based analysis to correct for the 48 multiple comparisons and giving a statistical significance level of 0.05/48 = 0.001.

## RESULTS

### Dose-finding experiments

In the initial, dose-finding cohort of mice, aged septic mice had reduced survival with increasing CS dose over the 24-hour timepoint and showed worse survival compared to young septic mice at CS doses above 1.6 mg/g (Supplemental Table 1). Over the 8-day timepoint, aged septic mice had reduced survival with increasing CS dose and worse survival compared to young septic mice at CS doses above 1.0 mg/g (Supplemental Table 2). At both timepoints, aged septic mice showed increasing illness severity with increasing CS dose (Supplemental Fig. 1B, D) while young septic mice required a CS dose above 1.0 mg/g to show increased illness severity (Supplemental Fig. 1A, C). Based on these findings, the 1.6 mg/g and 1.0 mg/g doses were used for subsequent experiments at the 24-hour and 8-day timepoints, respectively.

### Acute phase experiments

In mice that were euthanized 24 hours after injections, aged septic mice showed worse physiologic dysfunction, but no increase in bacterial burden or metabolism compared to young mice. Specifically, aged septic mice had increased illness severity at 4-hours, 8-hours, and 24-hours post CS injection, greater fall in body temperature at 4-hours post CS injection, and reduced weight loss compared to young mice at 24-hour post-CS injection (Fig. 1A-C). Although septic mice (young and aged) had increased bacterial burden compared to their respective control mice, no differences were observed between young and aged septic mice (Fig. 1D). Septic mice (young and aged) also had decreased plasma glucose compared to their respective controls (Supplemental Fig. 2A). However, young and aged septic mice did not differ with regards to plasma glucose, triglycerides or cholesterol (Supplemental Fig. 2A-C).

**Figure 1.**
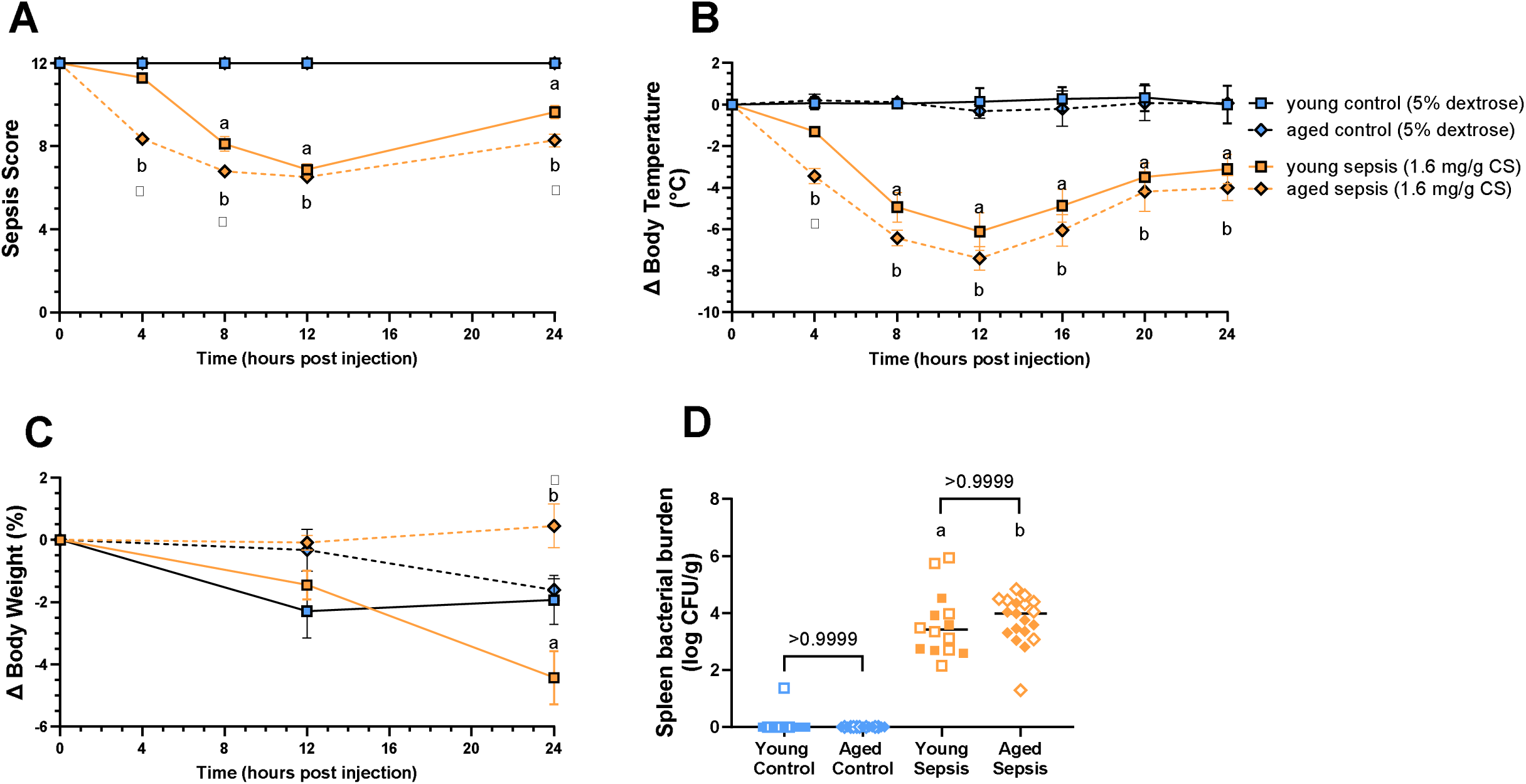
Severity of illness, change in body temperature, change in body weight, and spleen bacterial burden associated with a 24-hour septic insult [1.6 mg/g cecal slurry (CS)] in young and aged mice. Septic mice (young and aged) showed greater illness severity (A), loss of body temperature (B), and spleen bacterial counts (D) compared to control mice. Compared to young septic mice, aged septic mice also showed worse septic illness severity at 4-hours, 8-hours, and 24-hours post CS injection, lower body temperature at 4-hours post CS injection, and weight gain at 24-hours post CS injection. However, young and aged septic mice did not differ in spleen bacterial burden. Data were presented as mean +/− SEM (A-C) or as individual data points each representing an individual male (solid) or female (open) animal (D). Horizontal line indicates combined median for male and female animals. N = 3-25. [Statistical analysis: Two-way ANOVA + Tukey’s multiple comparisons test (A-C), Kruskal-Wallis test + Dunn’s multiple comparisons test (D).] ^a^p<0.05 vs Young Control; ^b^p<0.05 vs Aged Control; *p<0.05 Young Sepsis vs Aged Sepsis. Control = 5% dextrose, Sepsis = 1.6 mg/g CS.

At the 24-hour timepoint, aged septic mice also had elevated systemic inflammation compared to young septic mice. Specifically, aged septic mice had higher plasma concentrations of IL-6, TNF-α, IL-1β, and IFN-γ, as well as numerically higher CXCL1 compared to young septic mice (Fig. 2A-E). Although septic mice (both young and aged) had higher plasma IL-10 concentrations compared to their respective controls, no differences were observed between young and aged (Fig. 2F).

**Figure 2.**
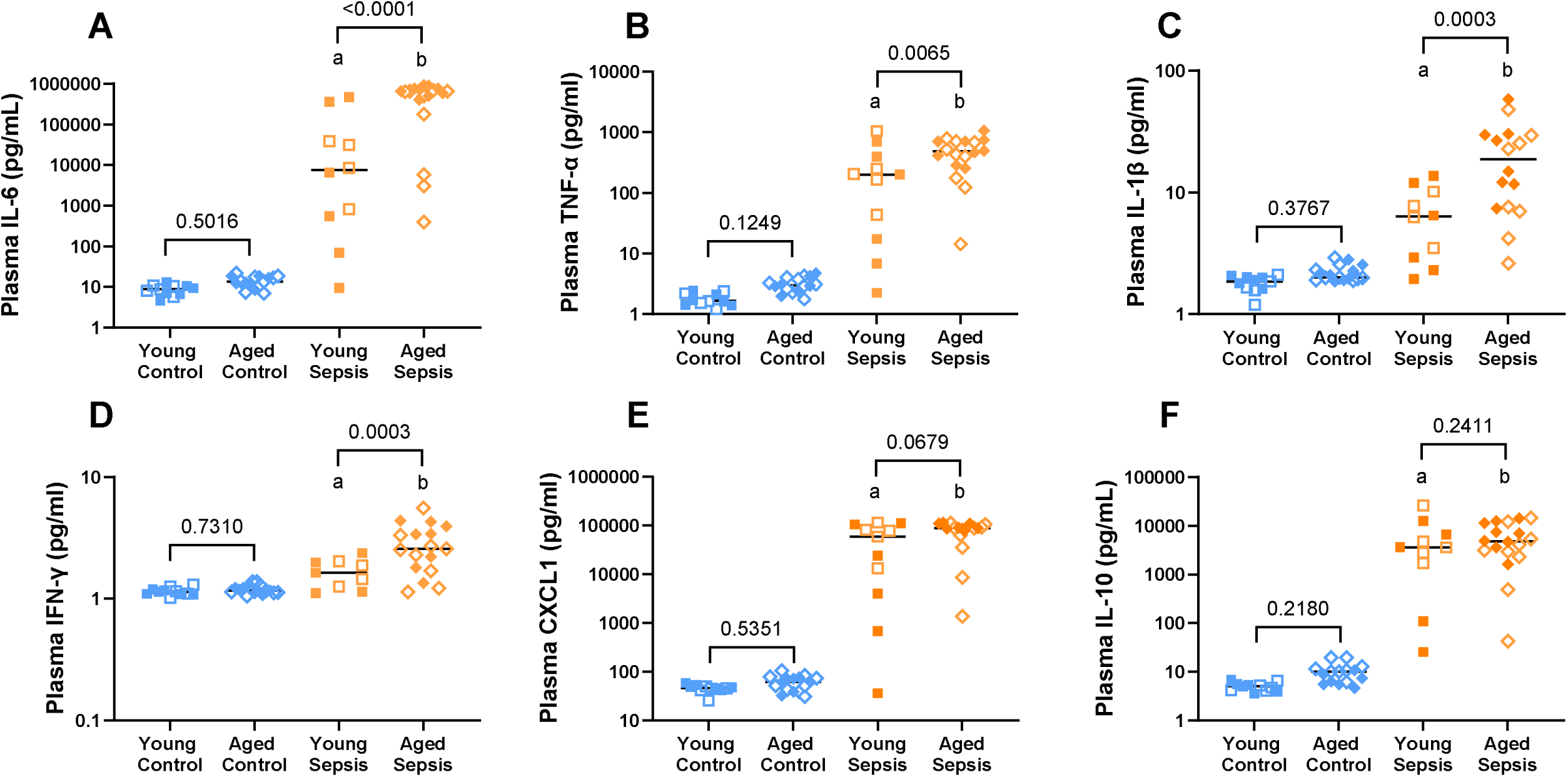
Systemic inflammation associated with 24-hour septic insult [cecal slurry (CS; 1.6mg/g)] in young and aged mice. Septic mice (both young and aged) had higher plasma concentrations of IL-6 (A), TNF-α (B), IL-1β (C), IFN-γ (D), CXCL1 (E), and IL-10 (F) compared to their respective controls. Aged septic mice also had significantly higher levels of circulating IL-6 (A), TNF-α (B), IL-1β (C), and IFN-γ (D) as well as numerically higher level of CXCL1 (E) compared to young septic mice. Plasma IL-10 (F) levels were not significantly different between young and aged septic mice. Each point represents an individual male (solid) or female (open) animal (A-F). Horizontal line indicates combined median for male and female animals. N = 3-18. [Statistical analysis: Two-way ANOVA + Uncorrected Fisher’s LSD test on log-transformed data (A-F).] ^a^p<0.05 vs Young Control; ^b^p<0.05 vs Aged Control. Control = 5% dextrose, Sepsis = 1.6 mg/g CS, IL-6 = interleukin-6, TNF-α = tumor necrosis factor alpha, IL-1β = interleukin-1 beta, IFN-γ = interferon gamma, CXCL1 = C-X-C motif chemokine ligand 1, IL-10 = interleukin-10.

In addition, aged septic mice had higher lung inflammation, but not lung edema during acute sepsis. Compared to young septic mice, aged septic mice had higher concentrations of BAL CXCL1, IL-6, TNF-α, and IL-1β, as well as numerically higher BAL IL-10 at 24-hours post CS injection (Fig. 3A-E). No differences were observed at this timepoint between age or treatment group for BAL IFN-γ concentrations, BAL inflammatory cell counts, BAL neutrophils (Fig. 3F-H), BAL protein concentrations, or lung wet-to-dry ratios (Supplemental Fig. 3A-B).

**Figure 3.**
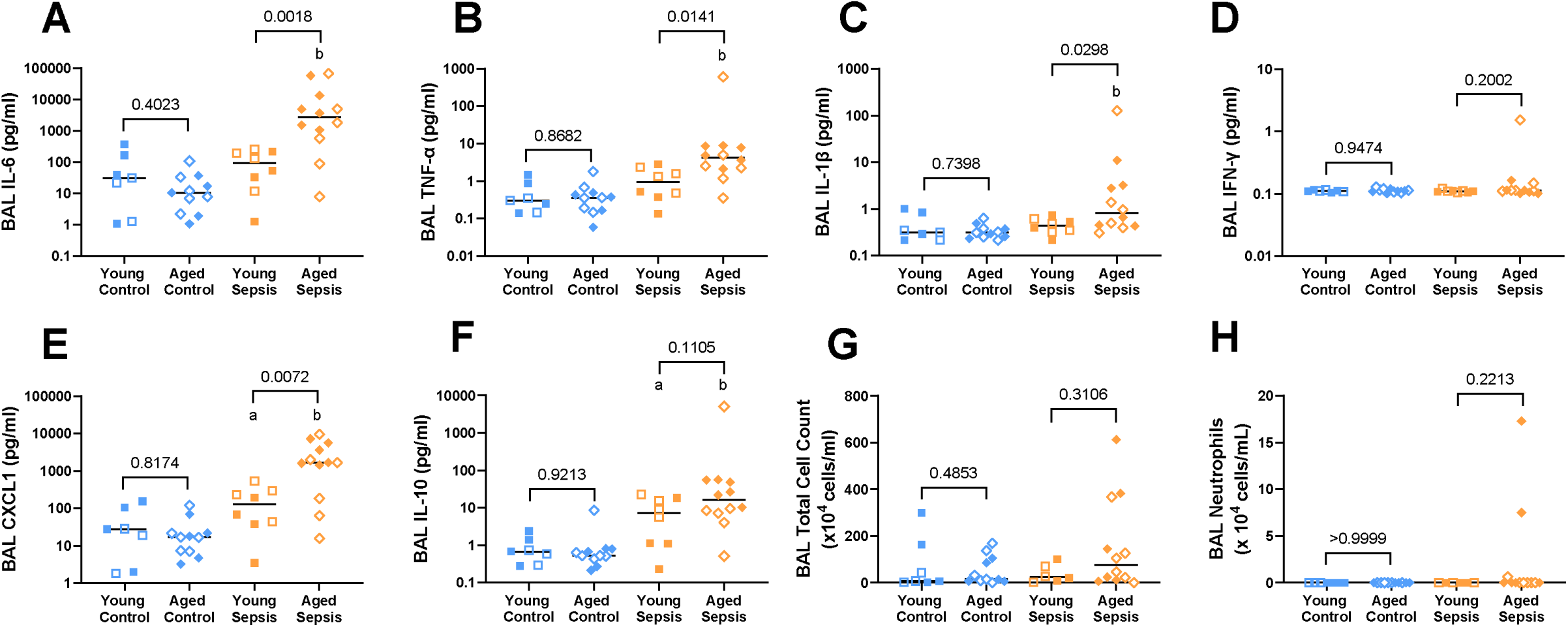
Lung inflammation associated with 24-hour septic insult [cecal slurry (CS; 1.6mg/g)] in young and aged mice. Aged septic mice had higher BAL concentrations of IL-6 (A), TNF-α (B), IL1-1β (C), CXCL1 (E), and IL-10 (F) compared to aged control mice. Young septic mice showed increases in BAL CXCL1 (E) and IL-10 (F) compared to young control mice. Compared to young septic mice, aged septic mice had higher BAL concentrations of IL-6 (A), TNF-α (B) IL1-1β (C), and CXCL1 (E) as well as numerically higher IL-10 (F). Sepsis and age had no effect on BAL IFN-γ concentrations (D), inflammatory cell counts (G), and number of neutrophils (H). Each point represents an individual male (solid circle) or female (open circle) animal (A-H). Horizontal line indicates combined median for male and female animals. N = 7-12. [Statistical analysis: Two-way ANOVA + Uncorrected Fisher’s LSD test on log-transformed data (A-G, I, J), Kruskal-Wallis test + Dunn’s multiple comparisons test (H).] ^a^p<0.05 vs Young Control; ^b^p<0.05 vs Aged Control. Control = 5% dextrose, Sepsis = 1.6 mg/g CS, BAL = bronchoalveolar lavage, IL-6 = interleukin-6, TNF-α = tumor necrosis factor alpha, IL-1β = interleukin-1 beta, IFN-γ = interferon gamma, CXCL1 = C-X-C motif chemokine ligand 1, IL-10 = interleukin-10.

Aged septic mice had increased liver inflammation and dysfunction, but not injury compared to young septic mice at 24 hours after CS injection. Specifically, aged septic mice had higher liver tissue mRNA expression of CXCL1, increased plasma total bilirubin concentrations, and reduced plasma albumin concentrations compared to young septic mice (Fig. 4A-C). No differences were observed between young and aged septic mice across any of the other cytokines measured via liver tissue mRNA expression or plasma concentrations of ALT and GGT (Supplemental Fig. 4A-G).

**Figure 4.**
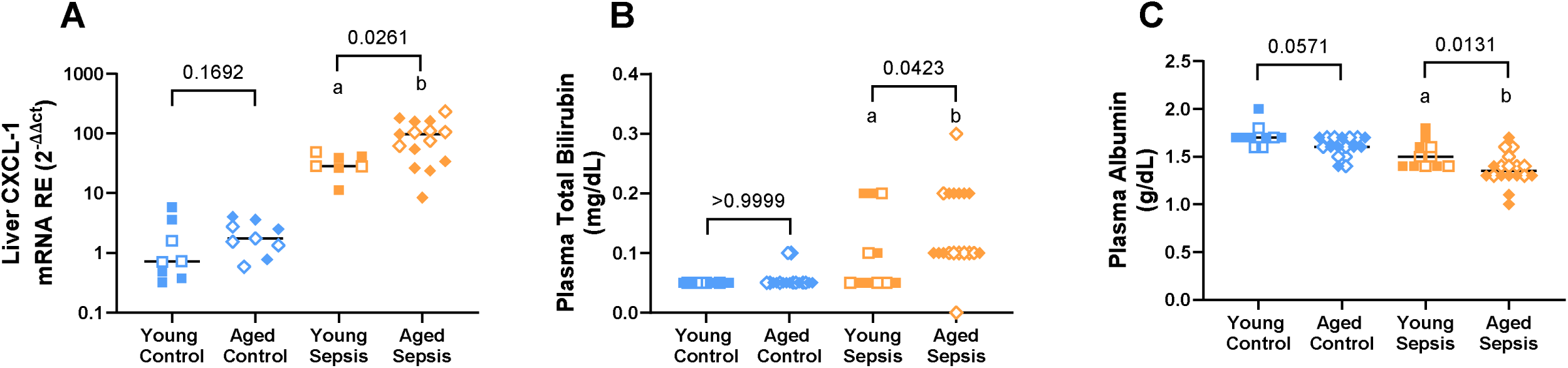
Liver inflammation and dysfunction associated with 24-hour septic insult [cecal slurry (CS; 1.6mg/g)] in young and aged mice. Septic mice (young and aged) septic mice had higher liver tissue mRNA expression of CXCL1 (A), increased plasma total bilirubin(B), and lower plasma albumin (C) compared to their respective control mice. Aged septic mice also had higher liver tissue mRNA expression of CXCL1 (A), increased plasma total bilirubin (B), and lower plasma albumin (C) compared to young septic mice. Each point represents an individual male (solid) or female (open) animal (A-C). Horizontal line indicates combined median for male and female animals. N = 7-18. [Statistical analysis: Two-way ANOVA + Uncorrected Fisher’s LSD test on log-transformed data (A-C)]. ^a^p<0.05 vs Young Control; ^b^p<0.05 vs Aged Control. Control = 5% dextrose, Sepsis = 1.6 mg/g CS, CXCL1 = C-X-C motif chemokine ligand.

Aged septic mice also had elevated kidney dysfunction and inflammation, but not injury during acute sepsis compared to young septic mice. At 24 hours after CS injection, aged septic mice had higher plasma urea nitrogen concentrations and kidney tissue mRNA expression of IL-6, IL-1β, and CXCL1 (Fig. 5A-D) compared to young septic mice. Despite numerical increases compared to aged control mice, aged septic mice did not differ compared to young septic mice with regards to kidney tissue mRNA expression of TNF-α, IFN-γ and IL-10, or kidney injury markers including KIM-1 and NGAL at both the mRNA and protein levels (Supplemental Fig. 5A-I).

**Figure 5.**
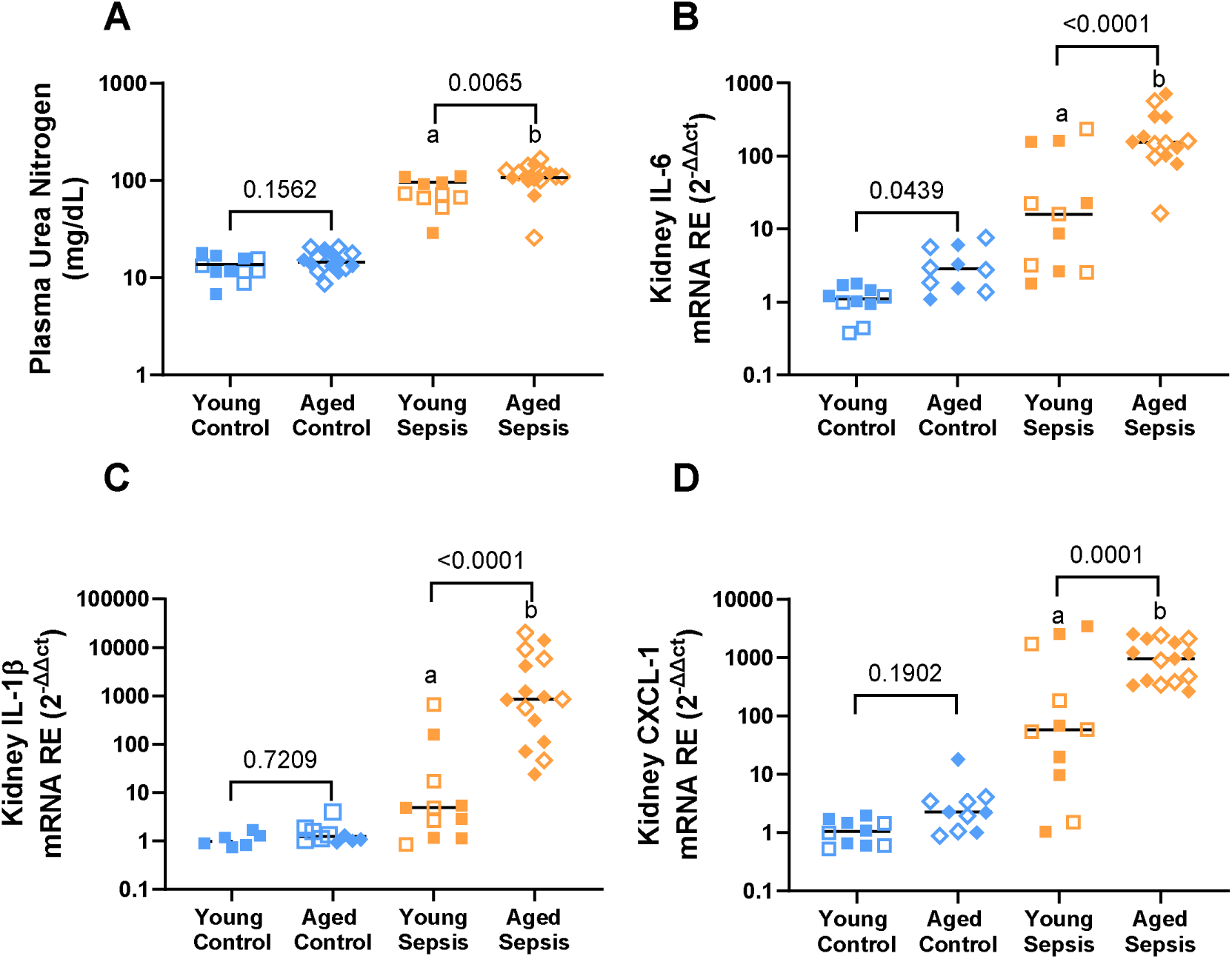
Kidney dysfunction and inflammation associated with 24-hour septic insult [cecal slurry (CS; 1.6mg/g)] in young and aged mice. Septic mice (young and aged) had elevated plasma urea nitrogen concentrations (A) as well as kidney tissue mRNA expression for IL-6 (B), IL-1β (C), and CXCL1 (D) compared to their respective control mice. Compared to young septic mice, aged septic mice had higher plasma urea nitrogen concentrations as well as kidney tissue mRNA expression forIL-6, IL-1β, and CXCL1. Each point represents an individual male (solid) or female (open) animal (A-D). Horizontal line indicates combined median for male and female animals. N = 10-15. [Statistical analysis: Two-way ANOVA + Uncorrected Fisher’s LSD test on log-transformed data (A-D)]. ^a^p<0.05 vs Young Control; ^b^p<0.05 vs Aged Control. Control = 5% dextrose, Sepsis = 1.6 mg/g CS, IL-6 = interleukin-6, IL-1β = interleukin-1 beta, CXCL1 = C-X-C motif chemokine ligand 1.

Lastly, aged septic mice had an increased endothelial injury and vascular permeability during acute sepsis compared to young septic mice. At 24-hours after CS injection, aged septic mice had higher plasma concentrations of soluble P-selectin and whole kidney fluorescent AngioSense accumulation compared to young septic mice (Fig. 6A-C). No differences were observed between young and aged septic mice for whole lung fluorescent AngioSense accumulation (Fig. 6D).

**Figure 6.**
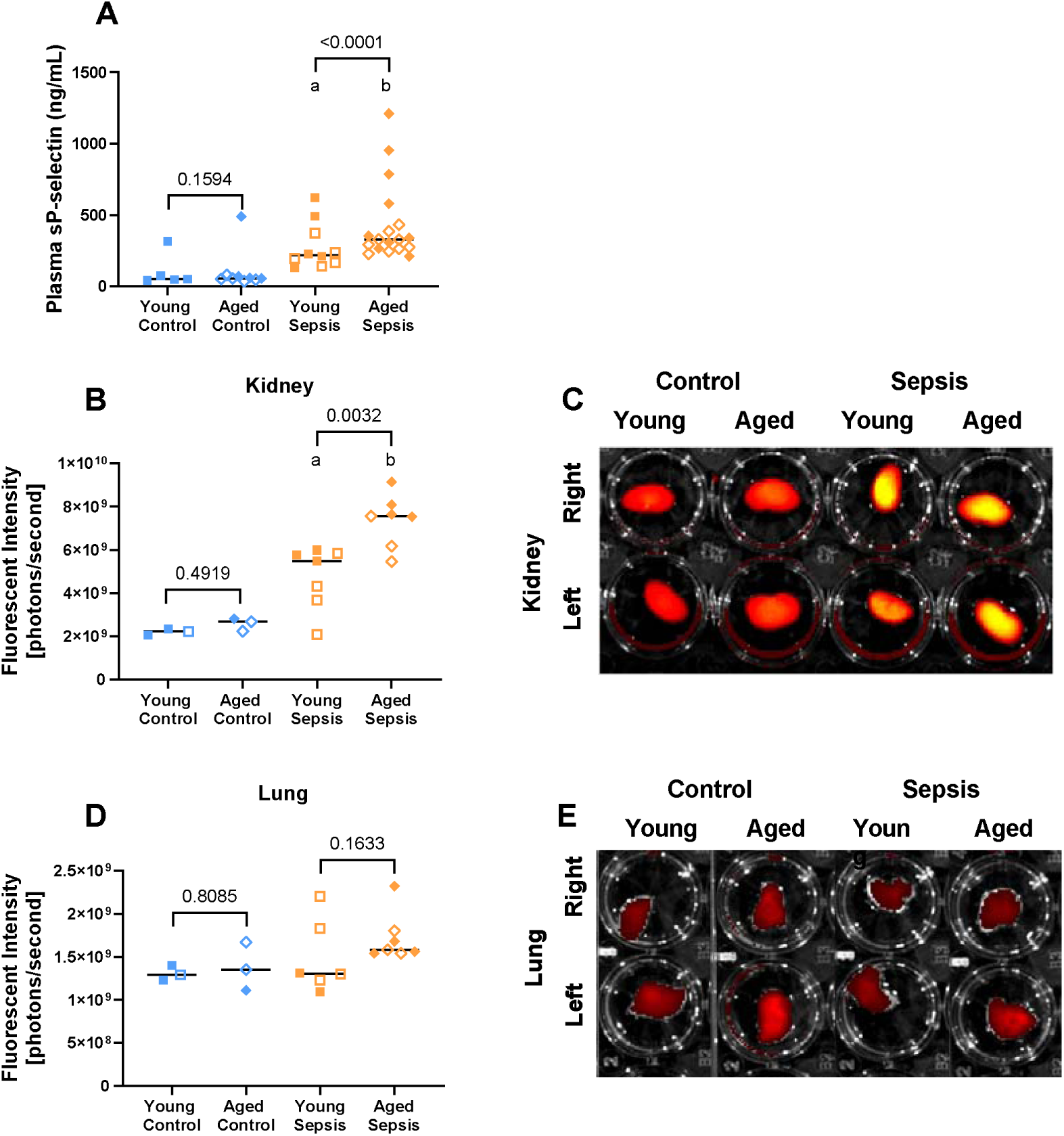
Endothelial injury and vascular permeability associated with 24-hour septic insult [cecal slurry (CS; 1.6mg/g)] in young and aged mice. Septic mice (both young and aged) had increased plasma concentrations of soluble p-selectin (A) and kidney accumulation of fluorescent AngioSense (B). Aged septic mice also had higher levels of plasma soluble p-selectin and kidney accumulation of fluorescent AngioSense, as well as numerically higher lung accumulation of fluorescent AngioSense (D) compared to young septic mice. Lung accumulation of fluorescent AngioSense was not different between septic and control mice for either age group. Representative images of kidneys (C) and lungs (E). Each point represents an individual male (solid circle) or female (open circle) animal (A, B, D). Horizontal line indicates combined median for male and female animals. N = 3-18. [Statistical analysis: Two-way ANOVA + Uncorrected Fisher’s LSD test on log-transformed data (A-F).] ^a^p<0.05 vs Young Control; ^b^p<0.05 vs Aged Control. Control = 5% dextrose, Sepsis = 1.6 mg/g cecal slurry.

### Recovery phase experiments

Compared to young septic mice, aged septic mice had worse physiologic dysfunction but not bacterial burden or metabolism after recovery from sepsis. Aged septic mice showed worse illness severity compared to young septic mice at all timepoints between 3-hours and 48-hours post CS injection, as well as greater loss of body temperature at 6-hour and 12-hours post CS injection (Fig. 7A-B). Aged septic mice had greater weight loss compared to control mice at all timepoints after 24-hours post CS injection (Fig. 7C). However, aged septic mice showed reduced weight loss at 24-hours post CS injection, yet greater weight loss at all timepoints after 72-hours post CS injection compared to young septic mice. At 8-days post CS injection, spleen bacterial counts, plasma glucose, plasma cholesterol, and plasma triglycerides were all similar between septic and control mice across either age group (Supplemental Fig. 6A-D).

**Figure 7.**
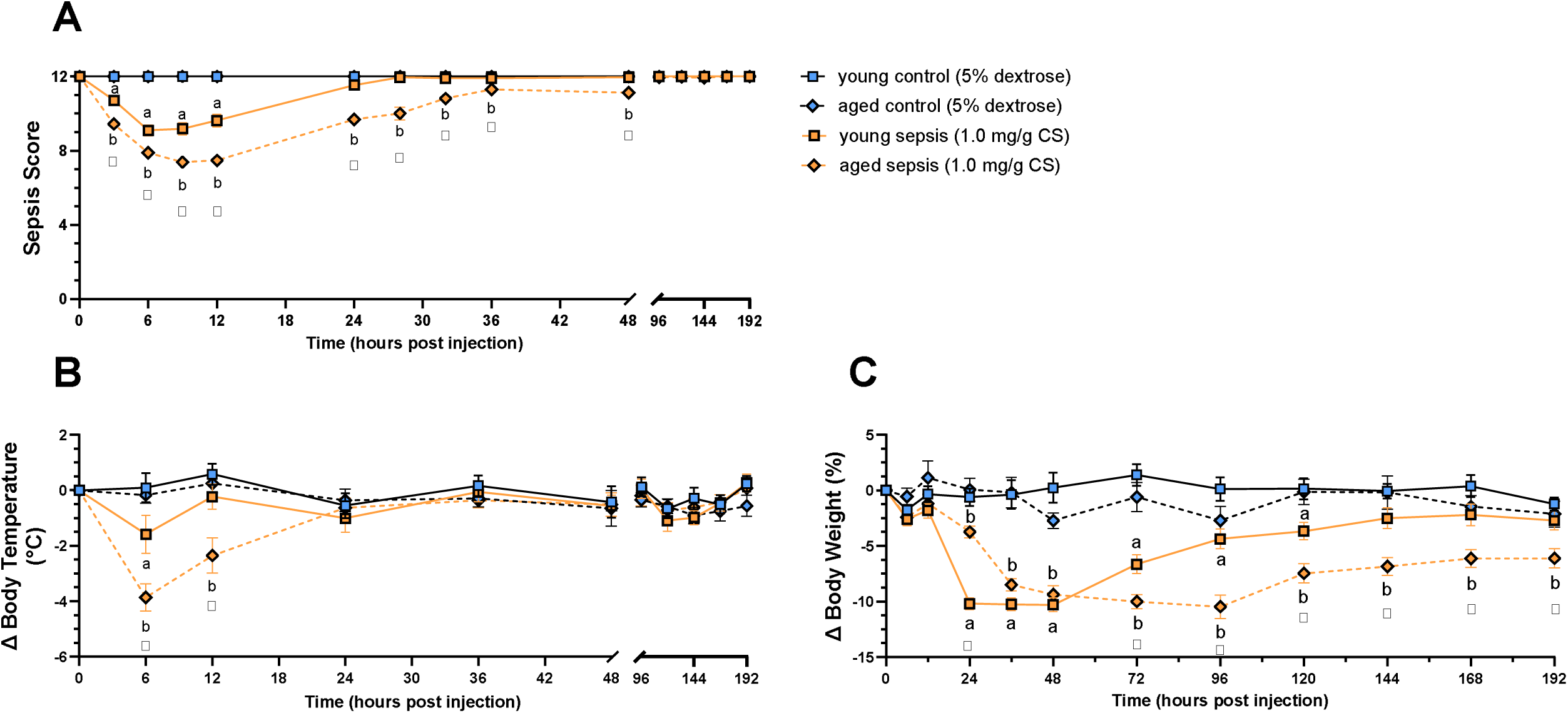
Severity of illness and change in body temperature and body weight associated with 8-day recovery from septic insult [1.0 mg/g cecal slurry (CS)] in young and aged mice. Septic mice (both young and aged) showed greater illness severity at 3, 6, 9, 12 hours after CS injection (A), loss of body temperature at 6-hours after CS injection (B), and weight loss at 24, 36, 48, 72, 96, 120 hours after CS injection (C) compared to control mice. In addition, aged septic mice showed greater illness severity at timepoints beyond 12 hours after CS injection, loss in body temperature at 12-hours after CS injection, and weight loss at timepoints beyond 120 hours after CS injection compared to aged controls. Compared to young septic mice, aged septic mice also had greater illness severity at 3, 6, 9, 12, 24, 28, 32, 36, and 48 hours after CS injection, loss of body temperature at 6 and 12 hours after CS injection, weight retention at 24-hours after CS injection, and weight loss at 72, 96, 120, 144, 168, and 192 hours after CS injection. Data were presented as combined mean for male and female animals +/− SEM (A-C). N = 8-19. [Statistical analysis: Two-way ANOVA + Tukey’s multiple comparisons test (A-C).] ^a^p<0.05 vs Young Control; ^b^p<0.05 vs Aged Control. Control = 5% dextrose, Sepsis = 1.0 mg/g cecal slurry.

At the 24-hour timepoint, aged septic mice had higher systemic inflammation compared to young septic mice. At 8-days post CS injection, only aged septic mice had higher plasma concentrations of inflammatory cytokines compared to control mice (Fig. 8). Compared to young septic mice, aged septic mice had higher plasma concentrations of TNF-α and IL-10, as well as numerically higher CXCL1 and IL-6 (Fig. 8A-D). Plasma concentrations of IL-1 β and IFN-γ were not different between treatment or age groups (Fig. 8E-F).

**Figure 8.**
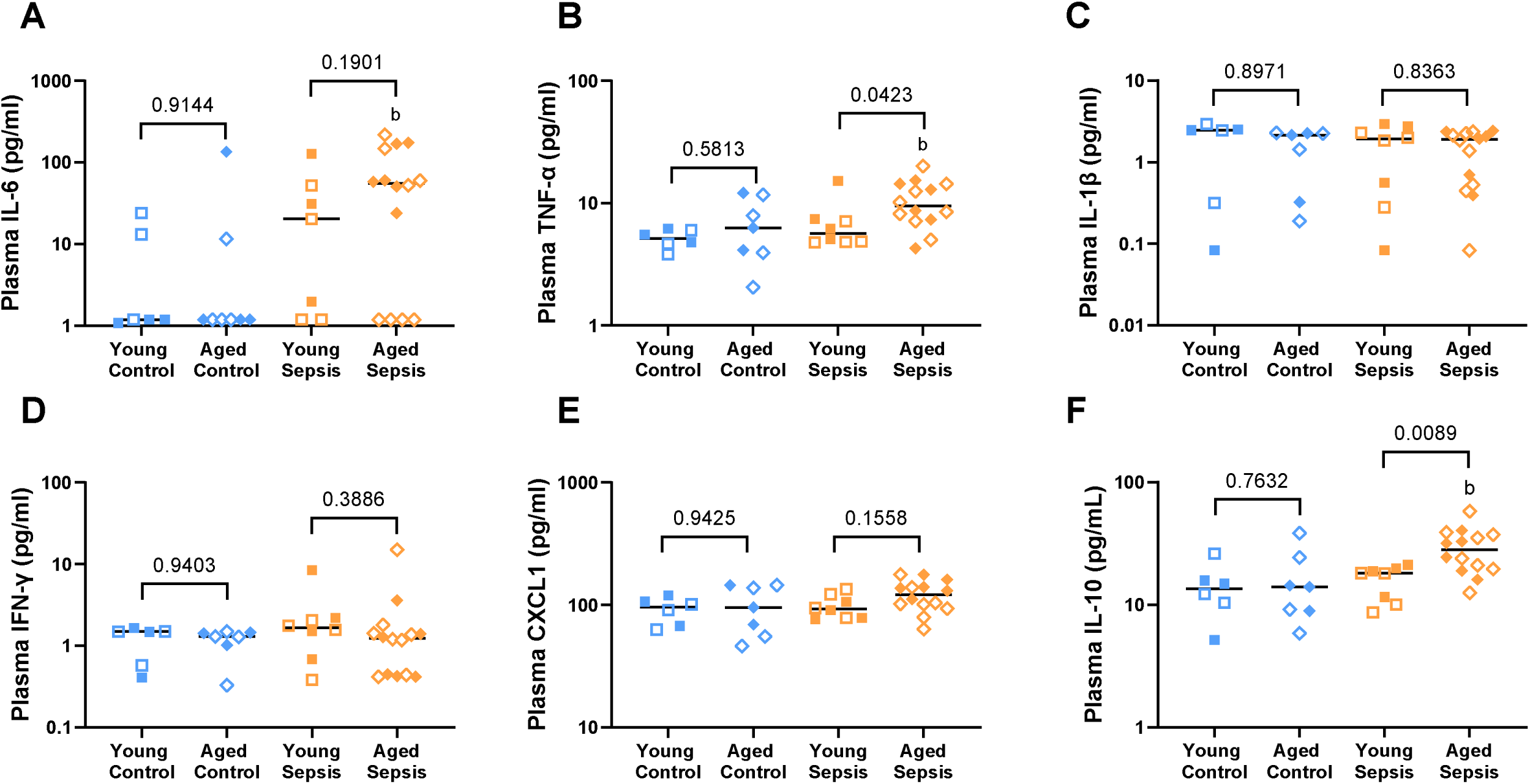
Systemic inflammation associated with 8-day recovery from septic insult [1.0 mg/g cecal slurry (CS)] in young and aged mice. Aged septic mice had increased plasma concentrations of IL-6 (A), TNF-α (B), and IL-10 (F), as well as numerically higher CXCL1 (p =0.0842; E) compared to aged control mice. However, no differences were observed across any of the pro-inflammatory cytokines measured between young septic mice and young control mice. Aged septic mice also showed higher plasma concentrations of TNF-α and IL-10, as well as numerically higher CXCL1 and IL-6 compared to young septic mice. Plasma concentrations of IL-1β (C), and IFN-γ (D) were not different across treatment or age groups. Each data point represents an individual male (solid circle) or female (open circle) animal (A-F). Horizontal line indicates combined median for male and female animals. N = 3-14. [Statistical analysis: Two-way ANOVA + Uncorrected Fisher’s LSD test on log-transformed data (A-F).] ^a^p<0.05 vs Young Control; ^b^p<0.05 vs Aged Control. Control = 5% dextrose, Sepsis = 1.0 mg/g cecal slurry, IL-6 = interleukin-6, TNF-α = tumor necrosis factor alpha, IL-1β = interleukin-1 beta, IFN-γ = interferon gamma, CXCL1 = C-X-C motif chemokine ligand 1, IL-10 = interleukin-10.

Aged septic mice did not have elevated lung inflammation or edema compared to young septic mice after recovery from sepsis. Specifically, no differences were observed between age groups across BAL cell counts, BAL differentials, any of the BAL cytokines, BAL neutrophils, BAL protein, or lung wet-to-dry ratios (Supplemental Fig. 7A-J). Similarly, aged septic mice did not differ compared to young septic mice across any markers of liver inflammation, dysfunction, or injury at this timepoint (Supplemental Fig.8A-F).

Aged septic mice did have increased kidney inflammation, dysfunction, and injury after recovery from sepsis. Specifically, aged septic mice had higher kidney tissue mRNA expression for CXCL1 and IL-10, as well as numerically higher IL-6, TNF-a, and IL-1β compared to young septic mice (Supplemental Fig. 9A-E). No differences were observed across treatment or age group for kidney tissue mRNA expression of IFN-γ (Supplemental Fig. 9F). Aged septic mice also had numerically higher plasma urea nitrogen and statistically higher kidney tissue expression of KIM-1 mRNA, KIM-1 protein, NGAL mRNA, as well as NGAL protein compared to young septic mice (Fig. 9A-G).

**Figure 9.**
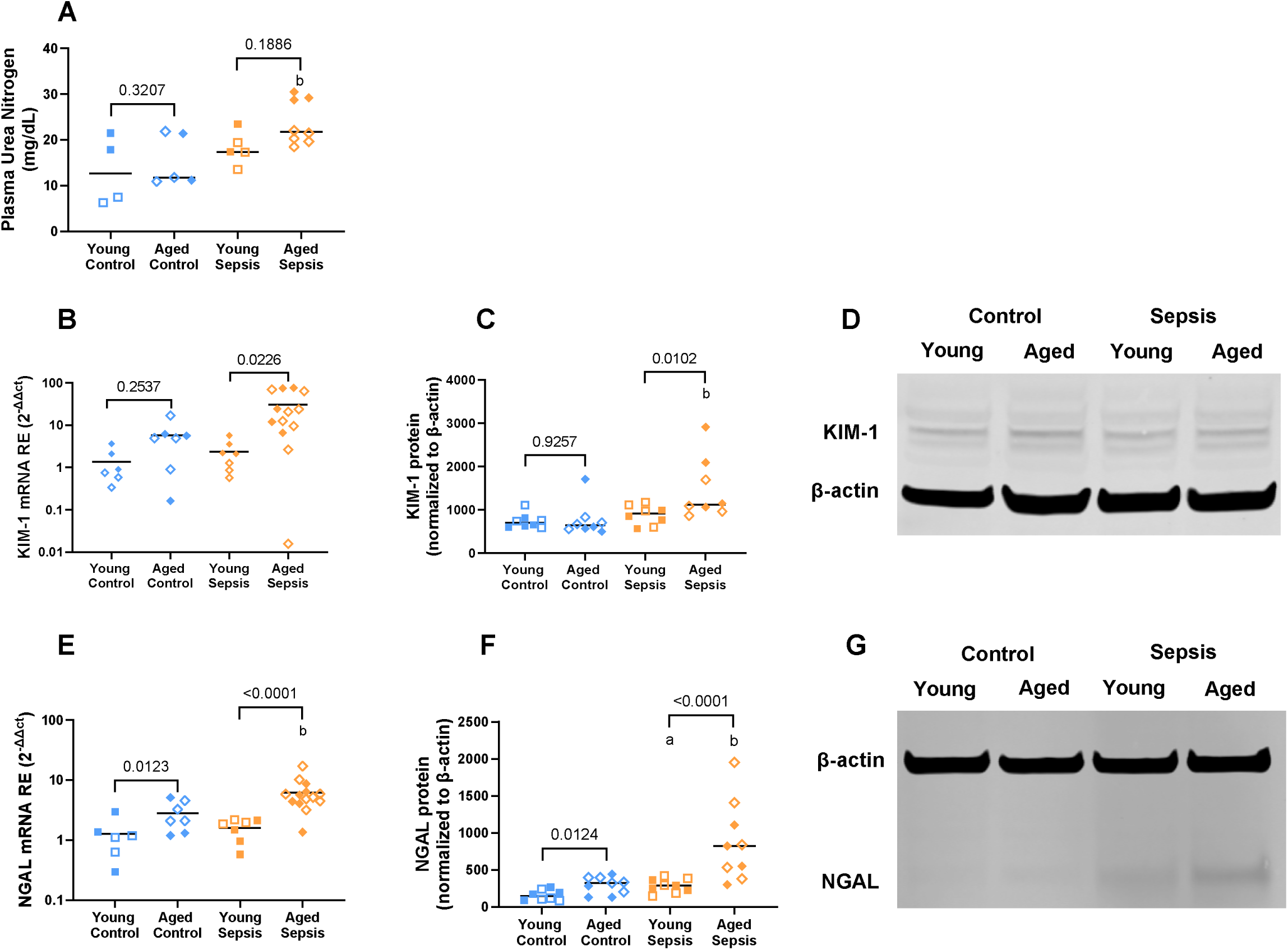
Kidney dysfunction and injury associated with 8-day recovery from septic insult [cecal slurry (CS; 1.0mg/g)] in young and aged mice. Compared to aged controls, aged septic mice had higher plasma urea nitrogen concentrations (A), kidney tissue expression of KIM-1 protein (C), NGAL mRNA (E), and NGAL protein (F), as well as numerically higher KIM-1 mRNA (p=0.0880; B). Aged septic mice also had higher kidney tissue expression of KIM-1 mRNA, KIM-1 protein, NGAL mRNA, and NGAL protein, as well as numerically higher plasma urea nitrogen (p=0.1886) compared to young septic mice. Representative images of KIM-1 Western blot (D) and NGAL Western blot (G). Each data point represents an individual male (solid circle) or female (open circle) animal (A-C, E-F). Horizontal line indicates the combined median of male and female animals. N = 4-14. [Statistical analysis: Two-way ANOVA + Uncorrected Fisher’s LSD test on log-transformed data (A-C, E-F)]. ^a^p<0.05 vs Young Control; ^b^p<0.05 vs Aged Control. Control = 5% dextrose, Sepsis = 1.0 mg/g cecal slurry, KIM-1 = kidney injury molecule-1, NGAL = neutrophil gelatinase associated lipocalin.

### Sex differences

Injection of CS into young or aged mice elicited a similar response across males and females for all outcomes at the 24-hour (Supplemental Table 3 and 4) and 8-day (Supplemental Table 5 and 6) timepoints.

## DISCUSSION

In this study, we tested the impact of older age on the pathophysiology of sepsis during both the acute and recovery phases, utilizing a polymicrobial abdominal sepsis model in 3- and 18-month-old male and female mice. We found that older age significantly worsened sepsis-induced physiologic dysfunction, systemic and organ-specific inflammation, endothelial injury, vascular permeability, and organ dysfunction during the acute phase. Additionally, older age prolonged sepsis-induced physiologic dysfunction, sustained systemic and organ-specific inflammation, and increased organ injury after recovery from sepsis. No biologically relevant differences between male and female mice were seen across different age or treatment groups at either timepoint. These results support our hypothesis that advanced age worsens outcomes and delays recovery in murine sepsis by dysregulating the immune response and increasing vascular endothelial permeability.

This study expands on previous research in lipopolysaccharide-(LPS)-, cecal ligation and puncture (CLP)-, and fecal slurry-induced models of sepsis ^22,51–53,63–65^ by investigating additional outcomes, including the impact of age on metabolism in sepsis, as reflected in blood glucose and lipid concentrations, and differential patterns of weight change during both the acute and recovery phases of sepsis. Notably, aged mice experienced less weight loss compared to young mice during the acute phase of sepsis but exhibited greater weight loss during the recovery phase of sepsis, which persisted throughout the course of recovery. This observation aligns with previous clinical studies on post-sepsis syndrome that have shown that elderly septic patients are more prone to persistent weight and muscle loss months after discharge from the intensive care unit ^66,67^. Thus, the similar pattern of sustained weight loss observed in aged septic mice might be attributed, at least in part, to muscle mass loss or a general reduction in nutrient absorption ^68^. In contrast, the weight gain observed in aged septic mice during the acute phase could indicate fluid retention, likely in the interstitial space, due to increased vascular leak. This finding supports previous research associating advanced age with increased vascular permeability in septic patients ^69–72^. Together, these findings highlight that the age-related changes in body weight and metabolism may be reflective of the more severe underlying pathophysiology of sepsis in elderly patients.

Mechanistically, dysregulated and persistent inflammation are core pathological features of severe sepsis, worsening disease progression by increasing endothelial vascular permeability ^34,73,74^. The abnormal release of pro-inflammatory cytokines triggers tissue-damaging immune cell infiltration, leading to endothelial injury and vascular leakage into organs ^75–77^. Aged septic mice, like severe sepsis patients, exhibited higher systemic and organ-specific levels of pro-inflammatory cytokines during acute sepsis, while the anti-inflammatory cytokine IL-10 was not elevated compared to young septic mice. This surge in pro-inflammatory cytokines such as IL-1β, IL-6, TNF-α, and IFN-γ suggests an increase in signals for innate immune cell recruitment, activation, and apoptosis in aged septic mice ^78^. Elevated CXCL1 and IL-1β in the plasma and organs of aged septic mice also suggest significant neutrophil involvement in this dysregulated response to acute sepsis ^78^. Even after recovery from sepsis, aged mice had persistently higher levels of TNF-α and IL-10, indicating ongoing inflammation and a compensatory anti-inflammatory response ^78^. This prolonged inflammation with high circulating levels of TNF-α, may also reflect an age-dependent exhaustion and depletion of adaptive immune cells post-recovery ^79,80^. These findings are consistent with clinical studies showing persistent inflammation in elderly patients after recovery from sepsis, which is associated with greater endothelial injury and vascular leakage ^38,46,47,51,53,65,81^. Thus, the more severe sepsis-induced organ injury and dysfunction observed in elderly patients may partly result from age-related immune dysregulation and increased endothelial vascular permeability.

Sepsis is characterized by organ injury leading to functional impairments in various organ systems ^82,83^. Importantly, dysfunction in the lungs, liver, and kidneys are major contributors to sepsis morbidity and mortality ^4,6,7,84–89^. We observed a modest increase in lung inflammation in aged septic mice, compared to young septic mice during the acute phase of sepsis. However, this inflammation did not persist in the recovery phase, nor did it disrupt the alveolar-capillary barrier at either timepoint, aligning with previous animal studies ^51,90^. Conversely, our study found that advanced age does exacerbate sepsis-induced liver and kidney dysfunction at the acute phase of sepsis. Notably, aged septic mice exhibited increased plasma levels of total bilirubin (liver) and urea nitrogen (kidney). These findings are consistent with well-documented symptoms of severe acute sepsis observed in elderly patients and aged rodent models, where liver and kidney dysfunction are prominent features ^7,22,44,63,65,91,92^. Our study builds on this previous work by investigating organ injury through tissue specific biomarkers at both the acute and recovery phases of sepsis. We found that aged septic mice exhibited increased kidney injury (KIM-1 and NGAL) after recovery from sepsis. Although previous studies have shown elevated markers of liver and kidney injury in aged mice with acute sepsis ^22,44,46,53,63,92^, our findings extend to the recovery period. These findings underscore the importance of evaluating outcomes across all phases of sepsis to better understand the unique vulnerabilities of aged individuals to sepsis-induced organ dysfunction and injury.

This study had several limitations. First, direct comparison of the acute and recovery phases of sepsis was not possible due to varying CS doses used for acute (1.6 mg/g) and recovery (1.0 mg/g) timepoints. These doses were chosen to achieve observable illness while allowing most septic mice to survive to the final timepoint. Second, we did not study the molecular mechanisms by which age exacerbates sepsis outcomes. Our data suggest increased vascular permeability in aged mice results from a dysregulated immune response and heightened endothelial injury. However, based on pervious findings we could speculate that advanced age also increases endothelial cell vulnerability to injury through apoptotic pathways ^70^, reduced antioxidant mechanisms ^45^, or structural changes ^93–97^. These possibilities will need to be tested in future studies. Lastly, the study was limited by its relatively mild illness severity and lack of robust organ injury, a limitation of murine sepsis studies. As mentioned previously, significant lung injury was not observed, aligning with our previous findings that CS alone, without a second pulmonary insult, does not induce robust lung injury despite high mortality rates ^98^.

Despite these limitations, the current study demonstrated that older age exacerbates sepsis-induced inflammation and endothelial vascular permeability in both sexes, leading to increased illness severity, multi-organ dysfunction, and mortality. Our findings also indicate that these adverse outcomes persist in aged mice beyond the acute phase of sepsis. This persistence results in chronic inflammation, sustained kidney injury, and increased mortality rates after recovery from sepsis. Consequently, our study highlights the need to reconsider the extrapolation of acute phase data derived from young, healthy male mice to the broader human population, particularly given that most septic patients are of an advanced age ^10–15,99^. Future sepsis research should prioritize the use of aged models in both sexes and extend measurements to include multi-organ outcomes over longer timepoints, to more accurately reflect the clinical reality and address the unique vulnerabilities of elderly patients. This approach will help in developing more effective therapeutic strategies tailored to improve outcomes for this high-risk population.

## ABBREVIATIONS

CS: cecal slurry
VUMC: Vanderbilt University Medical Center
NIA: National Institute on Aging
BAL: bronchoalveolar lavage
PBS: phosphate-buffered saline
CFU: colony forming units
IFN-γ: interferon gamma
IL-10: interleukin-10
IL-1β: interleukin-1 beta
IL-6: interleukin-6
CXCL1: C-X-C motif chemokine ligand 1
TNF-α: tumor necrosis factor alpha
GAPDH: glyceraldehyde-3-phosphate dehydrogenase
ALT: alanine aminotransferase
GGT: gamma-glutamyl transferase
KIM-1: kidney injury molecule-1
NGAL: neutrophil gelatinase associated lipocalin
ALT: alanine aminotransferase
GGT: gamma-glutamyl transferase
ROUT: robust regression and outlier removal
LPS: lipopolysaccharide
CLP: cecal ligation and puncture

## ACKNOWLEDGEMENTS

We thank the Translational Pathology Shared Resource supported by NCI/NIH Cancer Center Support Grant P30CA068485.

## SUPPLEMENTAL DIGITAL CONTENT

### Supplemental Tables

**Supplemental Table 1.** Comparison of survival rates (%) between young and aged mice 24-hours after injection of cecal slurry.

| Treatment | Young Survival, % [n] | Aged Survival, % [n] | Log-Rank P-Value |
| --- | --- | --- | --- |
| 1.0 mg/g CS | 100% [5] | 100% [8] | >0.9999 |
| 1.2 mg/g CS | - | 100% [6] |  |
| 1.4 mg/g CS | - | 60% [5] |  |
| 1.6 mg/g CS | 100% [17] | 97% [34] | 0.4795 |
| 2.0 mg/g CS | 100% [10] | 45% [11] | 0.0070 |

**Supplemental Table 2.**
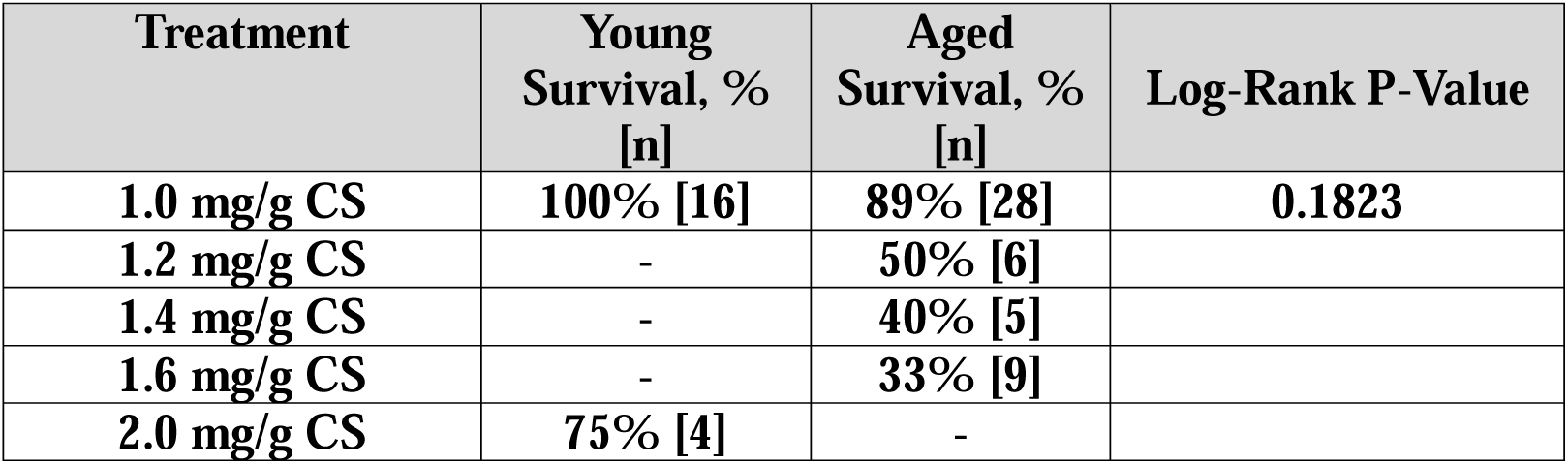
Comparison of survival rates (%) between young and aged mice 8-days after injection of cecal slurry.

| Treatment | Young Survival, % [n] | Aged Survival, % [n] | Log-Rank P-Value |
| --- | --- | --- | --- |
| 1.0 mg/g CS | 100% [16] | 89% [28] | 0.1823 |
| 1.2 mg/g CS | - | 50% [6] |  |
| 1.4 mg/g CS | - | 40% [5] |  |
| 1.6 mg/g CS | - | 33% [9] |  |
| 2.0 mg/g CS | 75% [4] | - |  |

**Supplemental Table 3.**
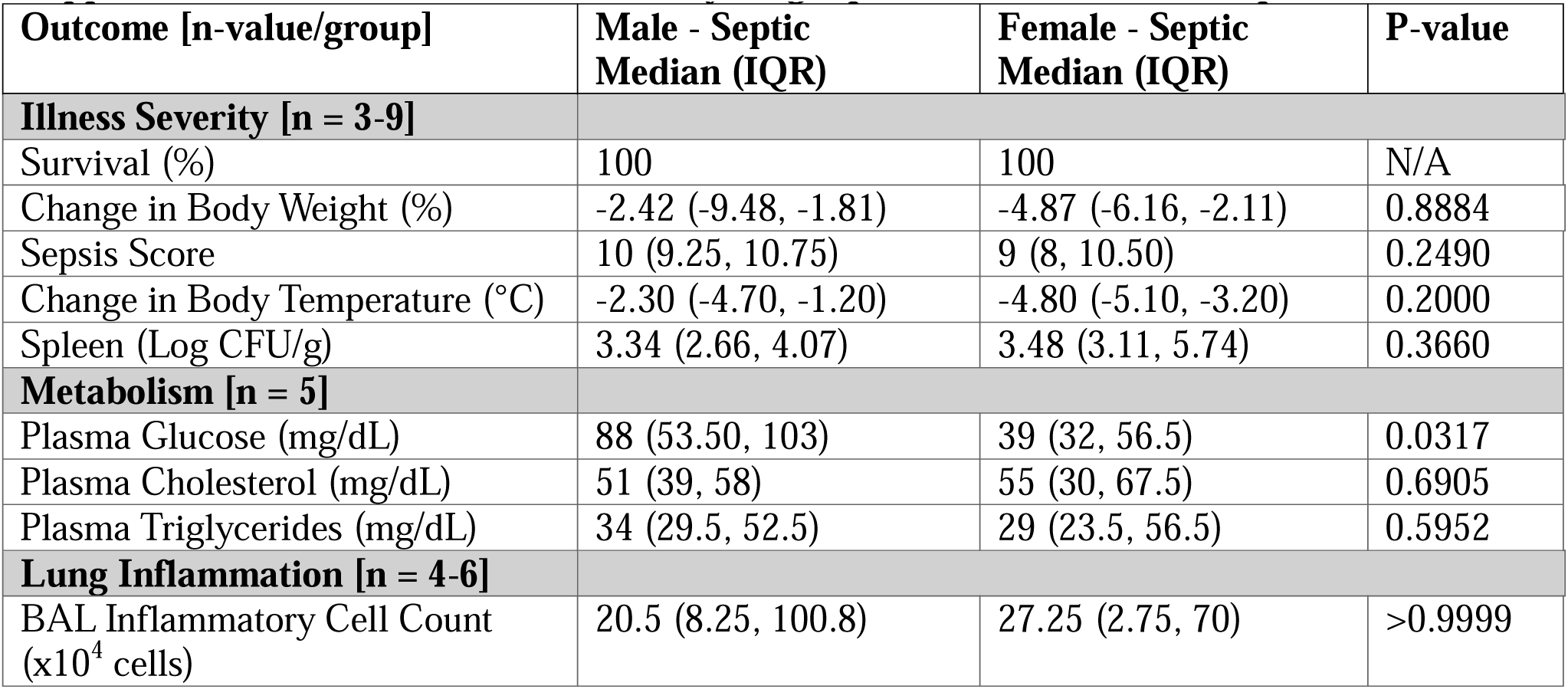
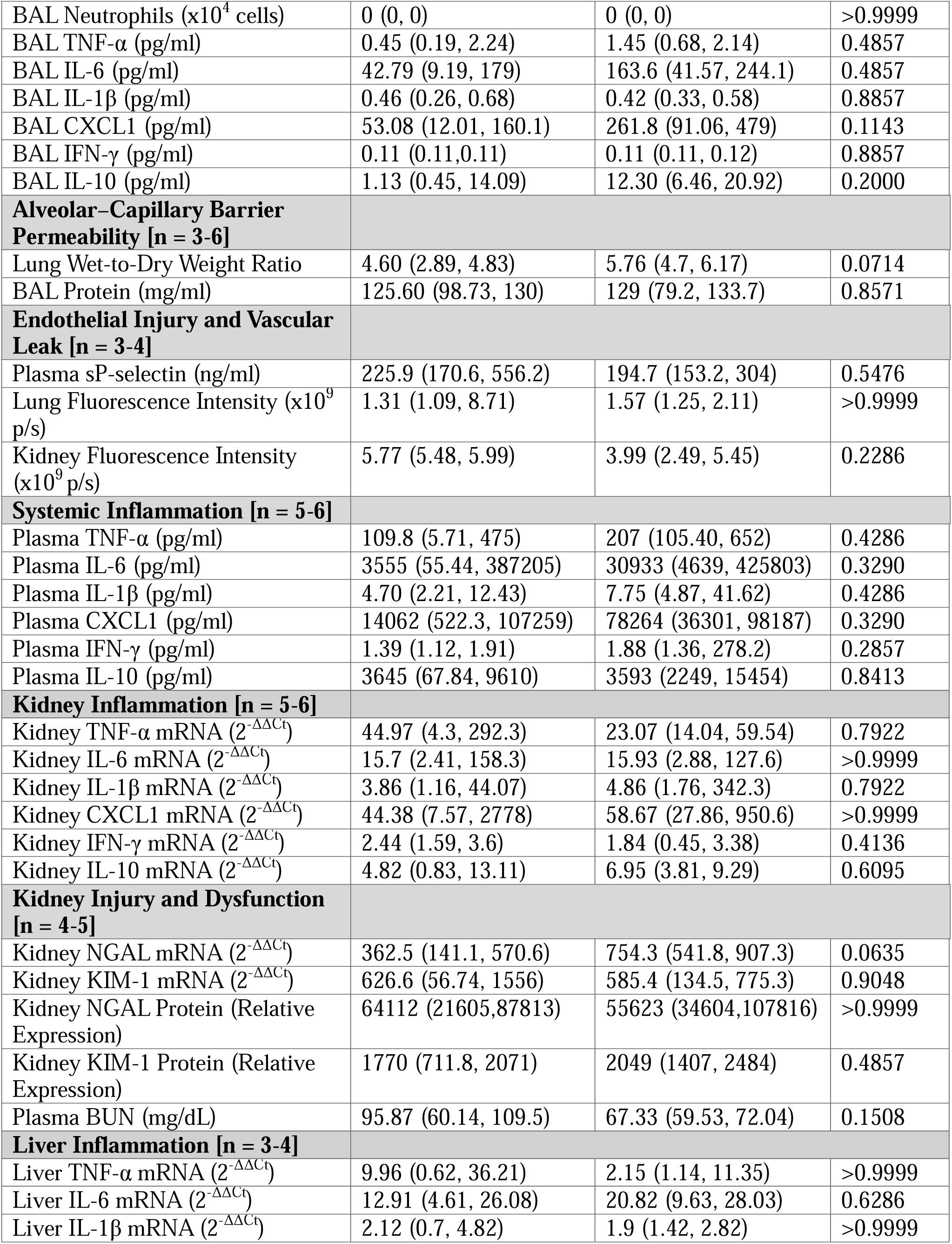
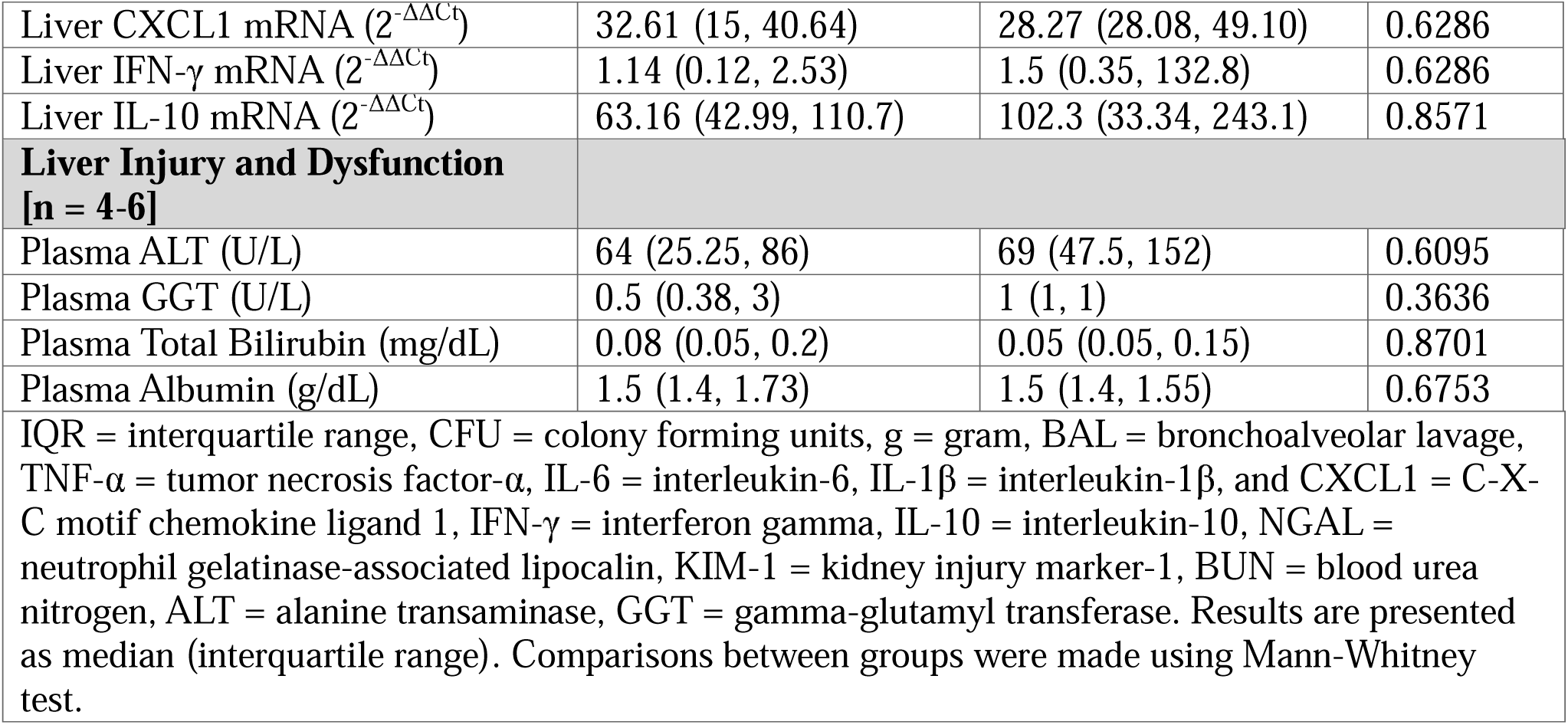
Sex differences for young septic mice at 24-hour timepoint.

| Outcome [n-value/group] | Male - Septic Median (IQR) | Female - Septic Median (IQR) | P-value |
| --- | --- | --- | --- |
| **Illness Severity [n = 3-9]** |  |  |  |
| Survival (%) | 100 | 100 | N/A |
| Change in Body Weight (%) | -2.42 (-9.48, -1.81) | -4.87 (-6.16, -2.11) | 0.8884 |
| Sepsis Score | 10 (9.25, 10.75) | 9 (8, 10.50) | 0.2490 |
| Change in Body Temperature (°C) | -2.30 (-4.70, -1.20) | -4.80 (-5.10, -3.20) | 0.2000 |
| Spleen (Log CFU/g) | 3.34 (2.66, 4.07) | 3.48 (3.11, 5.74) | 0.3660 |
| **Metabolism [n = 5]** |  |  |  |
| Plasma Glucose (mg/dL) | 88 (53.50, 103) | 39 (32, 56.5) | 0.0317 |
| Plasma Cholesterol (mg/dL) | 51 (39, 58) | 55 (30, 67.5) | 0.6905 |
| Plasma Triglycerides (mg/dL) | 34 (29.5, 52.5) | 29 (23.5, 56.5) | 0.5952 |
| **Lung Inflammation [n = 4-6]** |  |  |  |
| BAL Inflammatory Cell Count (x10<sup>4</sup> cells) | 20.5 (8.25, 100.8) | 27.25 (2.75, 70) | >0.9999 |
| BAL Neutrophils ($\times 10^4$ cells) | 0 (0, 0) | 0 (0, 0) | >0.9999 |
| BAL TNF-α (pg/ml) | 0.45 (0.19, 2.24) | 1.45 (0.68, 2.14) | 0.4857 |
| BAL IL-6 (pg/ml) | 42.79 (9.19, 179) | 163.6 (41.57, 244.1) | 0.4857 |
| BAL IL-1β (pg/ml) | 0.46 (0.26, 0.68) | 0.42 (0.33, 0.58) | 0.8857 |
| BAL CXCL1 (pg/ml) | 53.08 (12.01, 160.1) | 261.8 (91.06, 479) | 0.1143 |
| BAL IFN-γ (pg/ml) | 0.11 (0.11, 0.11) | 0.11 (0.11, 0.12) | 0.8857 |
| BAL IL-10 (pg/ml) | 1.13 (0.45, 14.09) | 12.30 (6.46, 20.92) | 0.2000 |
| **Alveolar–Capillary Barrier Permeability [n = 3-6]** |  |  |  |
| Lung Wet-to-Dry Weight Ratio | 4.60 (2.89, 4.83) | 5.76 (4.7, 6.17) | 0.0714 |
| BAL Protein (mg/ml) | 125.60 (98.73, 130) | 129 (79.2, 133.7) | 0.8571 |
| **Endothelial Injury and Vascular Leak [n = 3-4]** |  |  |  |
| Plasma sP-selectin (ng/ml) | 225.9 (170.6, 556.2) | 194.7 (153.2, 304) | 0.5476 |
| Lung Fluorescence Intensity ($\times 10^9$ p/s) | 1.31 (1.09, 8.71) | 1.57 (1.25, 2.11) | >0.9999 |
| Kidney Fluorescence Intensity ($\times 10^9$ p/s) | 5.77 (5.48, 5.99) | 3.99 (2.49, 5.45) | 0.2286 |
| **Systemic Inflammation [n = 5-6]** |  |  |  |
| Plasma TNF-α (pg/ml) | 109.8 (5.71, 475) | 207 (105.40, 652) | 0.4286 |
| Plasma IL-6 (pg/ml) | 3555 (55.44, 387205) | 30933 (4639, 425803) | 0.3290 |
| Plasma IL-1β (pg/ml) | 4.70 (2.21, 12.43) | 7.75 (4.87, 41.62) | 0.4286 |
| Plasma CXCL1 (pg/ml) | 14062 (522.3, 107259) | 78264 (36301, 98187) | 0.3290 |
| Plasma IFN-γ (pg/ml) | 1.39 (1.12, 1.91) | 1.88 (1.36, 278.2) | 0.2857 |
| Plasma IL-10 (pg/ml) | 3645 (67.84, 9610) | 3593 (2249, 15454) | 0.8413 |
| **Kidney Inflammation [n = 5-6]** |  |  |  |
| Kidney TNF-α mRNA ($2^{-\Delta\Delta C_t}$) | 44.97 (4.3, 292.3) | 23.07 (14.04, 59.54) | 0.7922 |
| Kidney IL-6 mRNA ($2^{-\Delta\Delta C_t}$) | 15.7 (2.41, 158.3) | 15.93 (2.88, 127.6) | >0.9999 |
| Kidney IL-1β mRNA ($2^{-\Delta\Delta C_t}$) | 3.86 (1.16, 44.07) | 4.86 (1.76, 342.3) | 0.7922 |
| Kidney CXCL1 mRNA ($2^{-\Delta\Delta C_t}$) | 44.38 (7.57, 2778) | 58.67 (27.86, 950.6) | >0.9999 |
| Kidney IFN-γ mRNA ($2^{-\Delta\Delta C_t}$) | 2.44 (1.59, 3.6) | 1.84 (0.45, 3.38) | 0.4136 |
| Kidney IL-10 mRNA ($2^{-\Delta\Delta C_t}$) | 4.82 (0.83, 13.11) | 6.95 (3.81, 9.29) | 0.6095 |
| **Kidney Injury and Dysfunction [n = 4-5]** |  |  |  |
| Kidney NGAL mRNA ($2^{-\Delta\Delta C_t}$) | 362.5 (141.1, 570.6) | 754.3 (541.8, 907.3) | 0.0635 |
| Kidney KIM-1 mRNA ($2^{-\Delta\Delta C_t}$) | 626.6 (56.74, 1556) | 585.4 (134.5, 775.3) | 0.9048 |
| Kidney NGAL Protein (Relative Expression) | 64112 (21605, 87813) | 55623 (34604, 107816) | >0.9999 |
| Kidney KIM-1 Protein (Relative Expression) | 1770 (711.8, 2071) | 2049 (1407, 2484) | 0.4857 |
| Plasma BUN (mg/dL) | 95.87 (60.14, 109.5) | 67.33 (59.53, 72.04) | 0.1508 |
| **Liver Inflammation [n = 3-4]** |  |  |  |
| Liver TNF-α mRNA ($2^{-\Delta\Delta C_t}$) | 9.96 (0.62, 36.21) | 2.15 (1.14, 11.35) | >0.9999 |
| Liver IL-6 mRNA ($2^{-\Delta\Delta C_t}$) | 12.91 (4.61, 26.08) | 20.82 (9.63, 28.03) | 0.6286 |
| Liver IL-1β mRNA ($2^{-\Delta\Delta C_t}$) | 2.12 (0.7, 4.82) | 1.9 (1.42, 2.82) | >0.9999 |
| Liver CXCL1 mRNA ($2^{-\Delta\Delta C_t}$) | 32.61 (15, 40.64) | 28.27 (28.08, 49.10) | 0.6286 |
| Liver IFN-γ mRNA ($2^{-\Delta\Delta C_t}$) | 1.14 (0.12, 2.53) | 1.5 (0.35, 132.8) | 0.6286 |
| Liver IL-10 mRNA ($2^{-\Delta\Delta C_t}$) | 63.16 (42.99, 110.7) | 102.3 (33.34, 243.1) | 0.8571 |
| **Liver Injury and Dysfunction**<br>**[n = 4-6]** |  |  |  |
| Plasma ALT (U/L) | 64 (25.25, 86) | 69 (47.5, 152) | 0.6095 |
| Plasma GGT (U/L) | 0.5 (0.38, 3) | 1 (1, 1) | 0.3636 |
| Plasma Total Bilirubin (mg/dL) | 0.08 (0.05, 0.2) | 0.05 (0.05, 0.15) | 0.8701 |
| Plasma Albumin (g/dL) | 1.5 (1.4, 1.73) | 1.5 (1.4, 1.55) | 0.6753 |
| IQR = interquartile range, CFU = colony forming units, g = gram, BAL = bronchoalveolar lavage, TNF-α = tumor necrosis factor-α, IL-6 = interleukin-6, IL-1β = interleukin-1β, and CXCL1 = C-X-C motif chemokine ligand 1, IFN-γ = interferon gamma, IL-10 = interleukin-10, NGAL = neutrophil gelatinase-associated lipocalin, KIM-1 = kidney injury marker-1, BUN = blood urea nitrogen, ALT = alanine transaminase, GGT = gamma-glutamyl transferase. Results are presented as median (interquartile range). Comparisons between groups were made using Mann-Whitney test. | | | |

**Supplemental Table 4.**
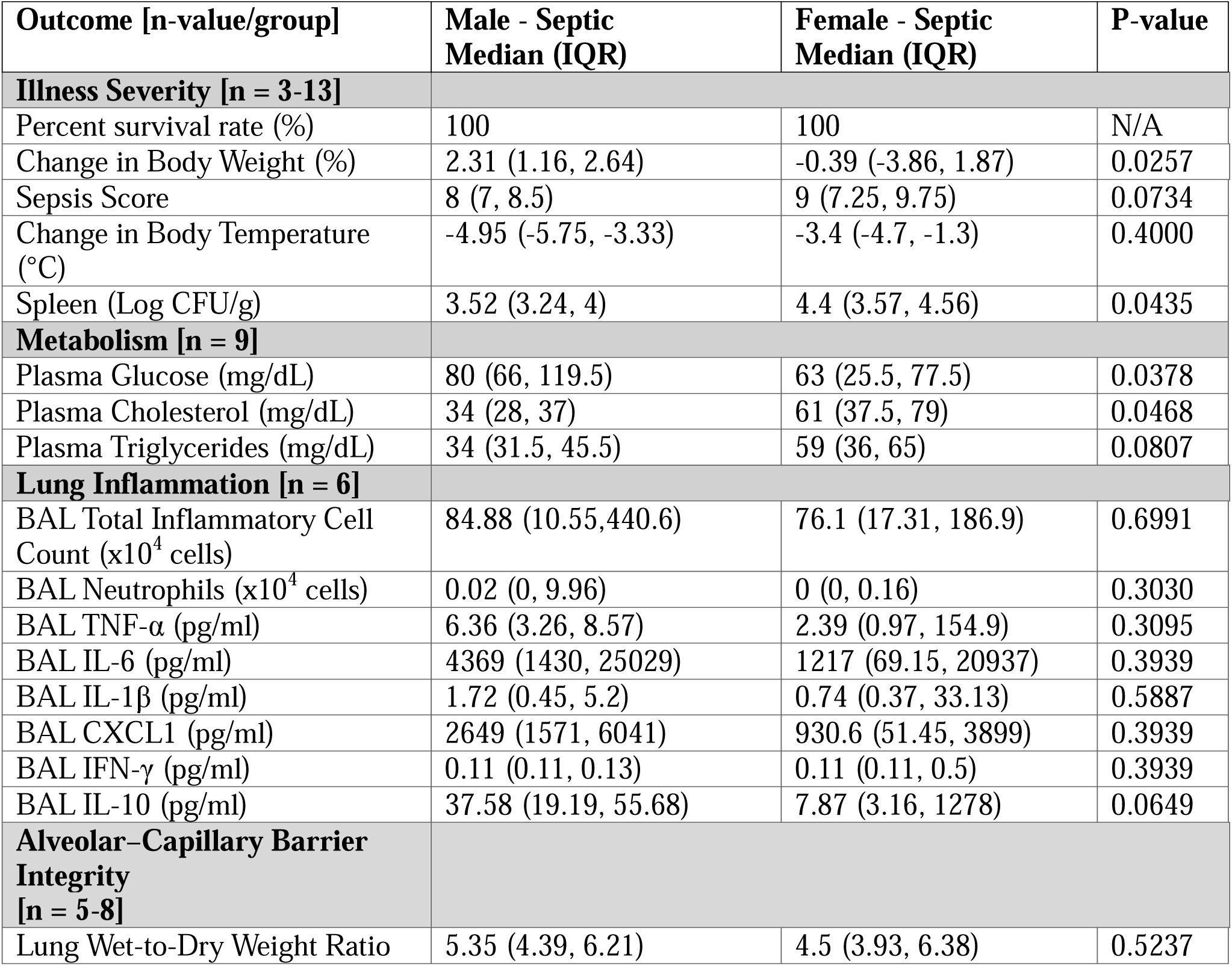
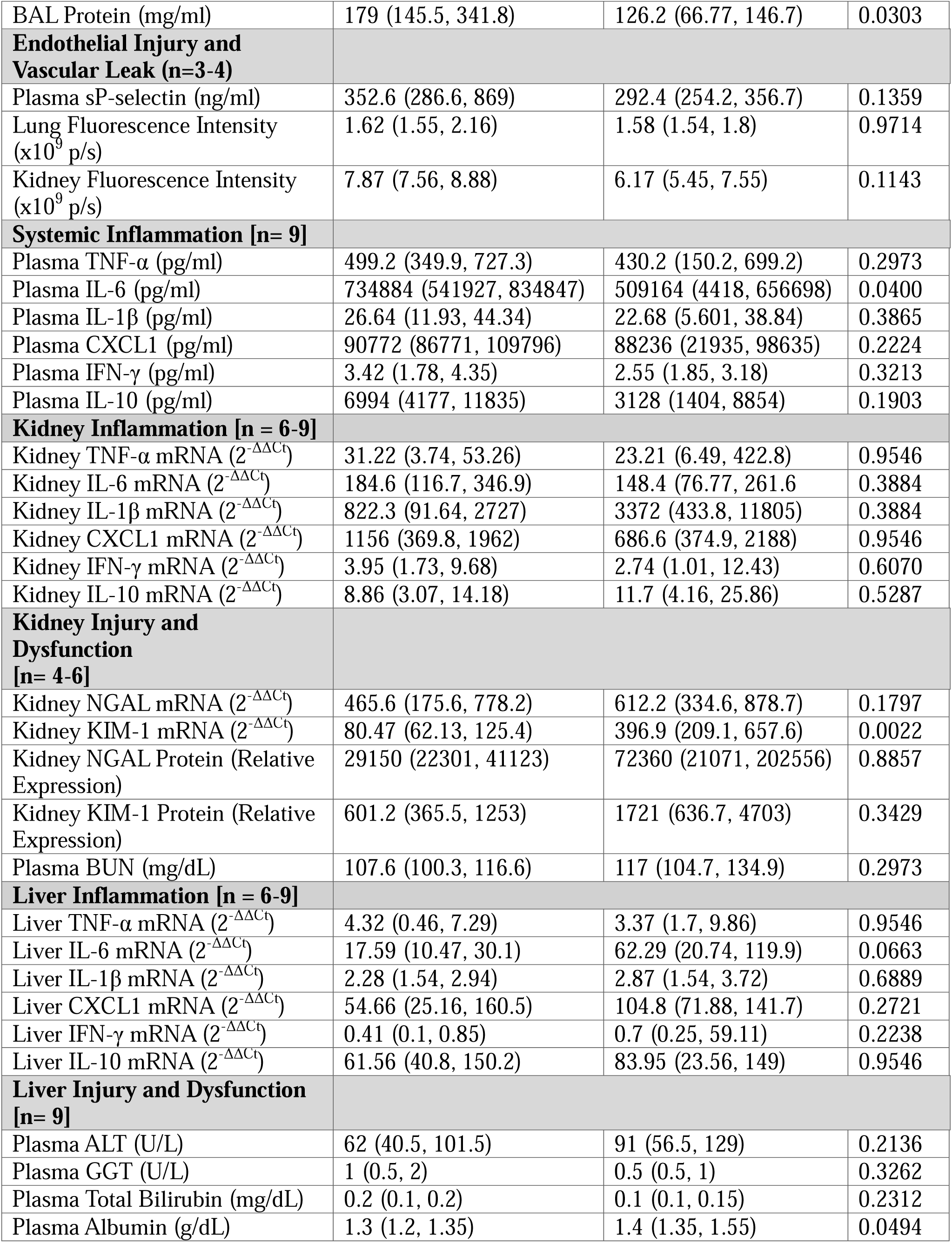
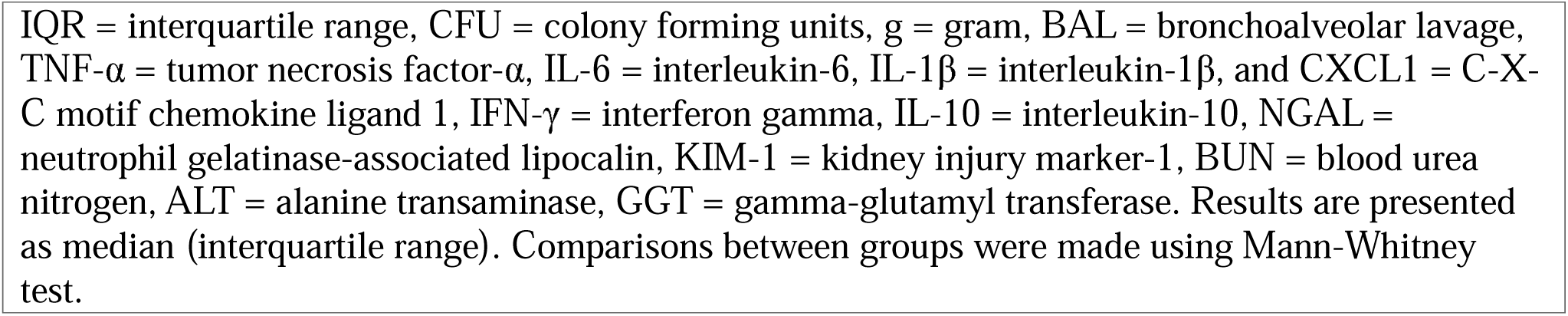
Sex differences for aged septic mice at 24-hour timepoint.

| Outcome [n-value/group] | Male - Septic<br>Median (IQR) | Female - Septic<br>Median (IQR) | P-value |
| --- | --- | --- | --- |
| **Illness Severity [n = 3-13]** |  |  |  |
| Percent survival rate (%) | 100 | 100 | N/A |
| Change in Body Weight (%) | 2.31 (1.16, 2.64) | -0.39 (-3.86, 1.87) | 0.0257 |
| Sepsis Score | 8 (7, 8.5) | 9 (7.25, 9.75) | 0.0734 |
| Change in Body Temperature (°C) | -4.95 (-5.75, -3.33) | -3.4 (-4.7, -1.3) | 0.4000 |
| Spleen (Log CFU/g) | 3.52 (3.24, 4) | 4.4 (3.57, 4.56) | 0.0435 |
| **Metabolism [n = 9]** |  |  |  |
| Plasma Glucose (mg/dL) | 80 (66, 119.5) | 63 (25.5, 77.5) | 0.0378 |
| Plasma Cholesterol (mg/dL) | 34 (28, 37) | 61 (37.5, 79) | 0.0468 |
| Plasma Triglycerides (mg/dL) | 34 (31.5, 45.5) | 59 (36, 65) | 0.0807 |
| **Lung Inflammation [n = 6]** |  |  |  |
| BAL Total Inflammatory Cell Count ($\times 10^4$ cells) | 84.88 (10.55, 440.6) | 76.1 (17.31, 186.9) | 0.6991 |
| BAL Neutrophils ($\times 10^4$ cells) | 0.02 (0, 9.96) | 0 (0, 0.16) | 0.3030 |
| BAL TNF-α (pg/ml) | 6.36 (3.26, 8.57) | 2.39 (0.97, 154.9) | 0.3095 |
| BAL IL-6 (pg/ml) | 4369 (1430, 25029) | 1217 (69.15, 20937) | 0.3939 |
| BAL IL-1β (pg/ml) | 1.72 (0.45, 5.2) | 0.74 (0.37, 33.13) | 0.5887 |
| BAL CXCL1 (pg/ml) | 2649 (1571, 6041) | 930.6 (51.45, 3899) | 0.3939 |
| BAL IFN-γ (pg/ml) | 0.11 (0.11, 0.13) | 0.11 (0.11, 0.5) | 0.3939 |
| BAL IL-10 (pg/ml) | 37.58 (19.19, 55.68) | 7.87 (3.16, 1278) | 0.0649 |
| **Alveolar–Capillary Barrier Integrity**<br>**[n = 5-8]** |  |  |  |
| Lung Wet-to-Dry Weight Ratio | 5.35 (4.39, 6.21) | 4.5 (3.93, 6.38) | 0.5237 |
| BAL Protein (mg/ml) | 179 (145.5, 341.8) | 126.2 (66.77, 146.7) | 0.0303 |
| **Endothelial Injury and Vascular Leak (n=3-4)** |  |  |  |
| Plasma sP-selectin (ng/ml) | 352.6 (286.6, 869) | 292.4 (254.2, 356.7) | 0.1359 |
| Lung Fluorescence Intensity ($\times 10^9$ p/s) | 1.62 (1.55, 2.16) | 1.58 (1.54, 1.8) | 0.9714 |
| Kidney Fluorescence Intensity ($\times 10^9$ p/s) | 7.87 (7.56, 8.88) | 6.17 (5.45, 7.55) | 0.1143 |
| **Systemic Inflammation [n= 9]** |  |  |  |
| Plasma TNF-α (pg/ml) | 499.2 (349.9, 727.3) | 430.2 (150.2, 699.2) | 0.2973 |
| Plasma IL-6 (pg/ml) | 734884 (541927, 834847) | 509164 (4418, 656698) | 0.0400 |
| Plasma IL-1β (pg/ml) | 26.64 (11.93, 44.34) | 22.68 (5.601, 38.84) | 0.3865 |
| Plasma CXCL1 (pg/ml) | 90772 (86771, 109796) | 88236 (21935, 98635) | 0.2224 |
| Plasma IFN-γ (pg/ml) | 3.42 (1.78, 4.35) | 2.55 (1.85, 3.18) | 0.3213 |
| Plasma IL-10 (pg/ml) | 6994 (4177, 11835) | 3128 (1404, 8854) | 0.1903 |
| **Kidney Inflammation [n = 6-9]** |  |  |  |
| Kidney TNF-α mRNA ($2^{-\Delta\Delta C_t}$) | 31.22 (3.74, 53.26) | 23.21 (6.49, 422.8) | 0.9546 |
| Kidney IL-6 mRNA ($2^{-\Delta\Delta C_t}$) | 184.6 (116.7, 346.9) | 148.4 (76.77, 261.6) | 0.3884 |
| Kidney IL-1β mRNA ($2^{-\Delta\Delta C_t}$) | 822.3 (91.64, 2727) | 3372 (433.8, 11805) | 0.3884 |
| Kidney CXCL1 mRNA ($2^{-\Delta\Delta C_t}$) | 1156 (369.8, 1962) | 686.6 (374.9, 2188) | 0.9546 |
| Kidney IFN-γ mRNA ($2^{-\Delta\Delta C_t}$) | 3.95 (1.73, 9.68) | 2.74 (1.01, 12.43) | 0.6070 |
| Kidney IL-10 mRNA ($2^{-\Delta\Delta C_t}$) | 8.86 (3.07, 14.18) | 11.7 (4.16, 25.86) | 0.5287 |
| **Kidney Injury and Dysfunction [n= 4-6]** |  |  |  |
| Kidney NGAL mRNA ($2^{-\Delta\Delta C_t}$) | 465.6 (175.6, 778.2) | 612.2 (334.6, 878.7) | 0.1797 |
| Kidney KIM-1 mRNA ($2^{-\Delta\Delta C_t}$) | 80.47 (62.13, 125.4) | 396.9 (209.1, 657.6) | 0.0022 |
| Kidney NGAL Protein (Relative Expression) | 29150 (22301, 41123) | 72360 (21071, 202556) | 0.8857 |
| Kidney KIM-1 Protein (Relative Expression) | 601.2 (365.5, 1253) | 1721 (636.7, 4703) | 0.3429 |
| Plasma BUN (mg/dL) | 107.6 (100.3, 116.6) | 117 (104.7, 134.9) | 0.2973 |
| **Liver Inflammation [n = 6-9]** |  |  |  |
| Liver TNF-α mRNA ($2^{-\Delta\Delta C_t}$) | 4.32 (0.46, 7.29) | 3.37 (1.7, 9.86) | 0.9546 |
| Liver IL-6 mRNA ($2^{-\Delta\Delta C_t}$) | 17.59 (10.47, 30.1) | 62.29 (20.74, 119.9) | 0.0663 |
| Liver IL-1β mRNA ($2^{-\Delta\Delta C_t}$) | 2.28 (1.54, 2.94) | 2.87 (1.54, 3.72) | 0.6889 |
| Liver CXCL1 mRNA ($2^{-\Delta\Delta C_t}$) | 54.66 (25.16, 160.5) | 104.8 (71.88, 141.7) | 0.2721 |
| Liver IFN-γ mRNA ($2^{-\Delta\Delta C_t}$) | 0.41 (0.1, 0.85) | 0.7 (0.25, 59.11) | 0.2238 |
| Liver IL-10 mRNA ($2^{-\Delta\Delta C_t}$) | 61.56 (40.8, 150.2) | 83.95 (23.56, 149) | 0.9546 |
| **Liver Injury and Dysfunction [n= 9]** |  |  |  |
| Plasma ALT (U/L) | 62 (40.5, 101.5) | 91 (56.5, 129) | 0.2136 |
| Plasma GGT (U/L) | 1 (0.5, 2) | 0.5 (0.5, 1) | 0.3262 |
| Plasma Total Bilirubin (mg/dL) | 0.2 (0.1, 0.2) | 0.1 (0.1, 0.15) | 0.2312 |
| Plasma Albumin (g/dL) | 1.3 (1.2, 1.35) | 1.4 (1.35, 1.55) | 0.0494 |

IQR = interquartile range, CFU = colony forming units, g = gram, BAL = bronchoalveolar lavage, TNF-α = tumor necrosis factor- $\alpha$ , IL-6 = interleukin-6, IL-1 $\beta$ = interleukin-1 $\beta$ , and CXCL1 = C-X-C motif chemokine ligand 1, IFN-γ = interferon gamma, IL-10 = interleukin-10, NGAL = neutrophil gelatinase-associated lipocalin, KIM-1 = kidney injury marker-1, BUN = blood urea nitrogen, ALT = alanine transaminase, GGT = gamma-glutamyl transferase. Results are presented as median (interquartile range). Comparisons between groups were made using Mann-Whitney test.

**Supplemental Table 5.** Sex differences for young mice at Day 8 timepoint.

| **Outcome [n-value/group]** | **Male - Septic**<br>**Median (IQR)** | **Female - Septic**<br>**Median (IQR)** | **P-value** |
| --- | --- | --- | --- |
| **Illness Severity [n = 4-5]** |  |  |  |
| Rate of Survival (%) | 100 | 100 | N/A |
| Change in Body Weight (%) | -2.88 (-5.82, -1.1) | -3.1 (-5.71, 1.74) | 0.7302 |
| Sepsis Score | 12 (12, 12) | 12 (12, 12) | >0.9999 |
| Change in Body Temperature (°C) | -0.4 (-0.6, -0.2) | 1.25 (0.75, 1.68) | 0.0159 |
| Spleen (Log CFU/g) | 0 (0, 0) | 0 (0, 0.52) | >0.9999 |
| **Metabolism [n = 4-5]** |  |  |  |
| Plasma Glucose (mg/dL) | 184 (165, 226) | 127.5 (111, 155.3) | 0.0317 |
| Plasma Cholesterol (mg/dL) | 41 (40.5, 42.5) | 34 (27, 36.5) | 0.0159 |
| Plasma Triglycerides (mg/dL) | 33 (28, 64) | 26 (24.5, 46.25) | 0.4524 |
| **Lung Inflammation [n = 4-5]** |  |  |  |
| BAL Total Inflammatory Cell Count (x10<sup>4</sup> cells) | 1.95 (1.13, 4.2) | 2.18 (0.79, 3) | 0.9365 |
| BAL Neutrophils (x10<sup>4</sup> cells) | 0 (0, 0) | 0 (0, 0) | >0.9999 |
| BAL TNF-α (pg/ml) | 6.01 (0.81, 6.82) | 4.83 (2.63, 6.55) | >0.9999 |
| BAL IL-6 (pg/ml) | 1.18 (0.81, 21.42) | 1.18 (0.59, 15.56) | 0.9048 |
| BAL IL-1β (pg/ml) | 0.48 (0.43, 0.52) | 0.39 (0.1, 0.46) | 0.1111 |
| BAL CXCL1 (pg/ml) | 90.59 (3.44, 109.8) | 86.46 (20.13, 124.7) | >0.9999 |
| BAL IFN-γ (pg/ml) | 0.14 (0.11, 0.18) | 0.11 (0.1, 0.13) | 0.1905 |
| BAL IL-10 (pg/ml) | 0.91 (0.68, 1.03) | 1.02 (0.55, 1.14) | 0.5556 |
| **Alveolar–Capillary Barrier Integrity [n = 4-5]** |  |  |  |
| Lung Wet-to-Dry Weight Ratio | 4.2 (3.17, 5.33) | 4.49 (3.59, 4.71) | 0.9048 |
| BAL Protein (mg/ml) | 172.7 (156.4, 242.9) | 131.3 (112.1, 143.8) | 0.0317 |
| **Systemic Inflammation [n= 4-5]** |  |  |  |
| Plasma TNF-α (pg/ml) | 6.21 (4.16, 11.28) | 4.83 (4.78, 6.55) | 0.4127 |
| Plasma IL-6 (pg/ml) | 11.9 (0.8, 79.35) | 10.77 (1.18, 44.41) | >0.9999 |
| Plasma IL-1β (pg/ml) | 2.02 (0.36, 2.84) | 1.92 (0.67, 2.22) | 0.7302 |
| Plasma CXCL1 (pg/ml) | 90.59 (67.45, 109.8) | 108.5 (82.36, 131.6) | 0.2857 |
| Plasma IFN-γ (pg/ml) | 1.52 (0.98, 5.33) | 1.67 (0.68, 2) | 0.9048 |
| Plasma IL-10 (pg/ml) | 18.77 (12.53, 20.54) | 14.06 (9, 18.16) | 0.2857 |
| **Kidney Inflammation [n= 5]** |  |  |  |
| Kidney TNF-α mRNA ($2^{-\Delta\Delta C_t}$) | 1.02 (0.55, 1.94) | 1.79 (1.17, 4.75) | 0.1508 |
| Kidney IL-6 mRNA ($2^{-\Delta\Delta C_t}$) | 1.18 (0.52, 2.5) | 1.85 (1.44, 3.99) | 0.5476 |
| Kidney IL-1β mRNA ($2^{-\Delta\Delta C_t}$) | 1.04 (0.65, 1.6) | 1.32 (1.03, 3.01) | 0.4206 |
| Kidney CXCL1 mRNA ($2^{-\Delta\Delta C_t}$) | 1.9 (0.4, 2.44) | 1.86 (0.85, 6.07) | 0.6905 |
| Kidney IFN-γ mRNA ($2^{-\Delta\Delta C_t}$) | 2.79 (1.56, 4.37) | 1.28 (0.81, 1.92) | 0.0952 |
| Kidney IL-10 mRNA ($2^{-\Delta\Delta C_t}$) | 1.23 (0.47, 3.2) | 1.96 (1.31, 7.02) | 0.4206 |
| **Kidney Injury and Dysfunction [n= 3-8]** |  |  |  |
| Kidney NGAL mRNA ($2^{-\Delta\Delta C_t}$) | 1.22 (0.67, 1.97) | 1.97 (1.87, 2.2) | 0.2286 |
| Kidney KIM-1 mRNA ($2^{-\Delta\Delta C_t}$) | 2.91 (2.13, 5.18) | 0.86 (0.57, 1.28) | 0.0571 |
| Kidney NGAL Protein (Relative Expression) | 256.5 (182, 344.8) | 339.5 (213.2, 410.7) | 0.3429 |
| Kidney KIM-1 Protein (Relative Expression) | 1149 (1066, 2093) | 958 (958, 1694) | 0.4000 |
| Plasma BUN (mg/dL) | 20.42 (17.38, 23.46) | 17.33 (13.55, 19.4) | 0.4000 |
| **Liver Inflammation [n= 4]** |  |  |  |
| Liver TNF-α mRNA ($2^{-\Delta\Delta C_t}$) | 1.64 (0.74, 2.53) | 1.33 (0.72, 1.6) | 0.6857 |
| Liver IL-6 mRNA ($2^{-\Delta\Delta C_t}$) | 2.61 (1, 4.17) | 1.39 (0.83, 2.52) | 0.4857 |
| Liver IL-1β mRNA ($2^{-\Delta\Delta C_t}$) | 1.36 (0.7, 2.08) | 0.96 (0.57, 1.26) | 0.6857 |
| Liver CXCL1 mRNA ($2^{-\Delta\Delta C_t}$) | 1.91 (1.23, 2.81) | 2.39 (1.28, 2.61) | 0.8857 |
| Liver IFN-γ mRNA ($2^{-\Delta\Delta C_t}$) | 0.36 (0.22, 0.48) | 0.36 (0.27, 0.67) | 0.8857 |
| Liver IL-10 mRNA ($2^{-\Delta\Delta C_t}$) | 2.41 (0.79, 4.42) | 1.38 (0.72, 2.1) | 0.6857 |
| **Liver Injury and Dysfunction [n= 4-5]** |  |  |  |
| Plasma ALT (U/L) | 14 (10, 51) | 13 (11, 18.75) | 0.6825 |
| Plasma GGT (U/L) | 0.5 (0.5, 0.5) | 0.5 (0.5, 0.5) | >0.9999 |
| Plasma Total Bilirubin (mg/dL) | 0.05 (0.05, 0.1) | 0.08 (0.05, 0.1) | >0.9999 |
| Plasma Albumin (g/dL) | 1.6 (1.6, 1.75) | 1.7 (1.55, 1.78) | 0.8016 |
| IQR = interquartile range, CFU = colony forming units, g = gram, BAL = bronchoalveolar lavage, TNF-α = tumor necrosis factor-α, IL-6 = interleukin-6, IL-1β = interleukin-1β, and CXCL1 = C-X-C motif chemokine ligand 1, IFN-γ = interferon gamma, IL-10 = interleukin-10, NGAL = neutrophil gelatinase-associated lipocalin, KIM-1 = kidney injury marker-1, BUN = blood urea nitrogen, ALT = alanine transaminase, GGT = gamma-glutamyl transferase. Results are presented as median (interquartile range). Comparisons between groups were made using Mann-Whitney test. | | | |

**Supplemental Table 6.** Sex differences for aged mice at Day 8 timepoint.

| Outcome [n-value/group] | Male - Septic Median (IQR) | Female - Septic Median (IQR) | P-value |
| --- | --- | --- | --- |
| **Illness Severity [n = 8-11]** |  |  |  |
| Rate of survival (%) | 88.89 | 100 | 0.2689 |
| Change in Body Weight (%) | -3.34 (-6.87, -0.5) | -5.62 (-7.91, -1.94) | 0.3282 |
| Sepsis Score | 12 (12, 12) | 12 (12, 12) | >0.9999 |
| Change in Body Temperature (°C) | 0 (-0.3, 0.75) | 0.15 (-0.23, 0.58) | 0.9004 |
| Spleen (Log CFU/g) | 0 (0, 0.4) | 0 (0, 0) | 0.4667 |
| **Metabolism [n = 6-8]** |  |  |  |
| Plasma Glucose (mg/dL) | 129 (99.5, 160.3) | 120 (97.75, 128) | 0.7002 |
| Plasma Cholesterol (mg/dL) | 38.5 (37, 47.75) | 30.5 (21.25, 34.5) | 0.0075 |
| Plasma Triglycerides (mg/dL) | 35 (29.75, 44.25) | 23.5 (21.25, 40.75) | 0.0881 |
| **Lung Inflammation [n = 6-11]** |  |  |  |
| BAL Total Inflammatory Cell Count (x10<sup>4</sup> cells) | 2.1 (1.54, 3.08) | 1.95 (1.39, 3.45) | 0.9382 |
| BAL Neutrophils (x10<sup>4</sup> cells) | 0 (0, 0) | 0 (0, 0) | >0.9999 |
| BAL TNF-α (pg/ml) | 7.04 (4.17, 14.03) | 6.6 (1.46, 11.9) | 0.5054 |
| BAL IL-6 (pg/ml) | 1.18 (0.58, 56.27) | 1.18 (0.72, 2.73) | >0.9999 |
| BAL IL-1β (pg/ml) | 0.47 (0.15, 0.93) | 0.38 (0.15, 0.5) | 0.6454 |
| BAL CXCL1 (pg/ml) | 96.49 (3.49, 130.7) | 71.38 (1.61, 129.7) | 0.5054 |
| BAL IFN-γ (pg/ml) | 0.12 (0.11, 0.13) | 0.13 (0.12, 0.16) | 0.2786 |
| BAL IL-10 (pg/ml) | 1.26 (0.78, 1.7) | 0.85 (0.42, 0.94) | 0.1605 |
| **Alveolar–Capillary Barrier Integrity [n = 8-11]** |  |  |  |
| Lung Wet-to-Dry Weight Ratio | 4.94 (4.44, 5.89) | 4.61 (3.96, 4.96) | 0.3953 |
| BAL Protein (mg/ml) | 166.4 (158.7, 186.9) | 159.1 (130.4, 164.5) | 0.2345 |
| **Systemic Inflammation [n= 6-8]** |  |  |  |
| Plasma TNF-α (pg/ml) | 8.11 (6.75, 14.03) | 9.36 (7.39, 13.89) | 0.7984 |
| Plasma IL-6 (pg/ml) | 54.48 (6.85, 142.3) | 26.9 (1.18, 126.3) | 0.6131 |
| Plasma IL-1β (pg/ml) | 2.02 (1.01, 2.35) | 1.63 (0.47, 2.25) | 0.3823 |
| Plasma CXCL1 (pg/ml) | 133.8 (108.3, 156) | 101.5 (82.79, 130.4) | 0.1304 |
| Plasma IFN-γ (pg/ml) | 1.34 (0.43, 1.6) | 1.3 (0.62, 1.71) | >0.9999 |
| Plasma IL-10 (pg/ml) | 22.73 (16.81, 32.63) | 29.56 (20, 38.62) | 0.4418 |
| **Kidney Inflammation [n= 8]** |  |  |  |
| Kidney TNF-α mRNA (2<sup>-$\Delta\Delta C_t$</sup>) | 1.25 (1.02, 3.65) | 3.08 (2.37, 6.25) | 0.1304 |
| Kidney IL-6 mRNA (2<sup>-$\Delta\Delta C_t$</sup>) | 1.86 (0.97, 4.88) | 3.62 (2.8, 7.96) | 0.1049 |
| Kidney IL-1β mRNA (2<sup>-$\Delta\Delta C_t$</sup>) | 1.27 (0.68, 3.05) | 2.63 (2.07, 5.27) | 0.0830 |
| Kidney CXCL1 mRNA (2<sup>-$\Delta\Delta C_t$</sup>) | 4.33 (1.36, 7.02) | 4.82 (3.04, 22.73) | 0.4567 |
| Kidney IFN-γ mRNA (2<sup>-$\Delta\Delta C_t$</sup>) | 1.07 (0.84, 1.22) | 1.36 (0.85, 2.99) | 0.2786 |
| Kidney IL-10 mRNA (2<sup>-$\Delta\Delta C_t$</sup>) | 2.55 (1.36, 7.11) | 9.54 (5.45, 17.11) | 0.0830 |
| **Kidney Injury and Dysfunction [n = 3-8]** |  |  |  |
| Kidney NGAL mRNA (2<sup>-$\Delta\Delta C_t$</sup>) | 5.04 (3.38, 6.98) | 5.52 (4.55, 9.1) | 0.4908 |
| Kidney KIM-1 mRNA (2<sup>-$\Delta\Delta C_t$</sup>) | 49.37 (10.56, 117.4) | 16.63 (4.37, 53.26) | 0.1812 |
| Kidney NGAL Protein (Relative Expression) | 821 (297.4, 1106) | 1126 (495.5, 1818) | 0.4000 |
| Kidney KIM-1 Protein (Relative Expression) | 1149 (1066, 2093) | 958 (861.2, 1694) | 0.4000 |
| Plasma BUN (mg/dL) | 29.21 (28.73, 30.46) | 20.31 (19.06, 21.77) | 0.0357 |
| **Liver Inflammation [n= 6]** |  |  |  |
| Liver TNF-α mRNA (2<sup>-$\Delta\Delta C_t$</sup>) | 1.63 (0.35, 2.77) | 2.08 (1.46, 2.74) | 0.5887 |
| Liver IL-6 mRNA (2<sup>-$\Delta\Delta C_t$</sup>) | 2.22 (0.43, 4.35) | 3.59 (3, 4.3) | 0.5887 |
| Liver IL-1β mRNA ($2^{-\Delta\Delta C_t}$) | 1.31 (0.31, 2.65) | 1.7 (1.44, 2.2) | 0.8182 |
| Liver CXCL1 mRNA ($2^{-\Delta\Delta C_t}$) | 2.79 (1.37, 5.69) | 1.14 (0.76, 2.51) | 0.1320 |
| Liver IFN-γ mRNA ($2^{-\Delta\Delta C_t}$) | 0.73 (0.42, 0.88) | 0.4 (0.3, 36.5) | 0.5887 |
| Liver IL-10 mRNA ($2^{-\Delta\Delta C_t}$) | 2.42 (0.3, 5.58) | 3.66 (3.01, 5.46) | 0.6991 |
| **Liver Injury and Dysfunction [n = 6-8]** |  |  |  |
| Plasma ALT (U/L) | 20 (10, 32.25) | 19 (9.25, 34.5) | 0.9366 |
| Plasma GGT (U/L) | 0.5 (0.5, 0.5) | 0.5 (0.5, 0.5) | >0.9999 |
| Plasma Total Bilirubin (mg/dL) | 0.1 (0.05, 0.18) | 0.05 (0.05, 0.1) | 0.3016 |
| Plasma Albumin (g/dL) | 1.65 (1.5, 1.7) | 1.45 (1.3, 1.58) | 0.1425 |
| IQR = interquartile range, CFU = colony forming units, g = gram, BAL = bronchoalveolar lavage, TNF-α = tumor necrosis factor-α, IL-6 = interleukin-6, IL-1β = interleukin-1β, and CXCL1 = C-X-C motif chemokine ligand 1, IFN-γ = interferon gamma, IL-10 = interleukin-10, NGAL = neutrophil gelatinase-associated lipocalin, KIM-1 = kidney injury marker-1, BUN = blood urea nitrogen, ALT = alanine transaminase, GGT = gamma-glutamyl transferase. Results are presented as median (interquartile range). Comparisons between groups were made using Mann-Whitney test. |  |  |  |

### Supplemental Figures

**Supplemental Figure 1.**
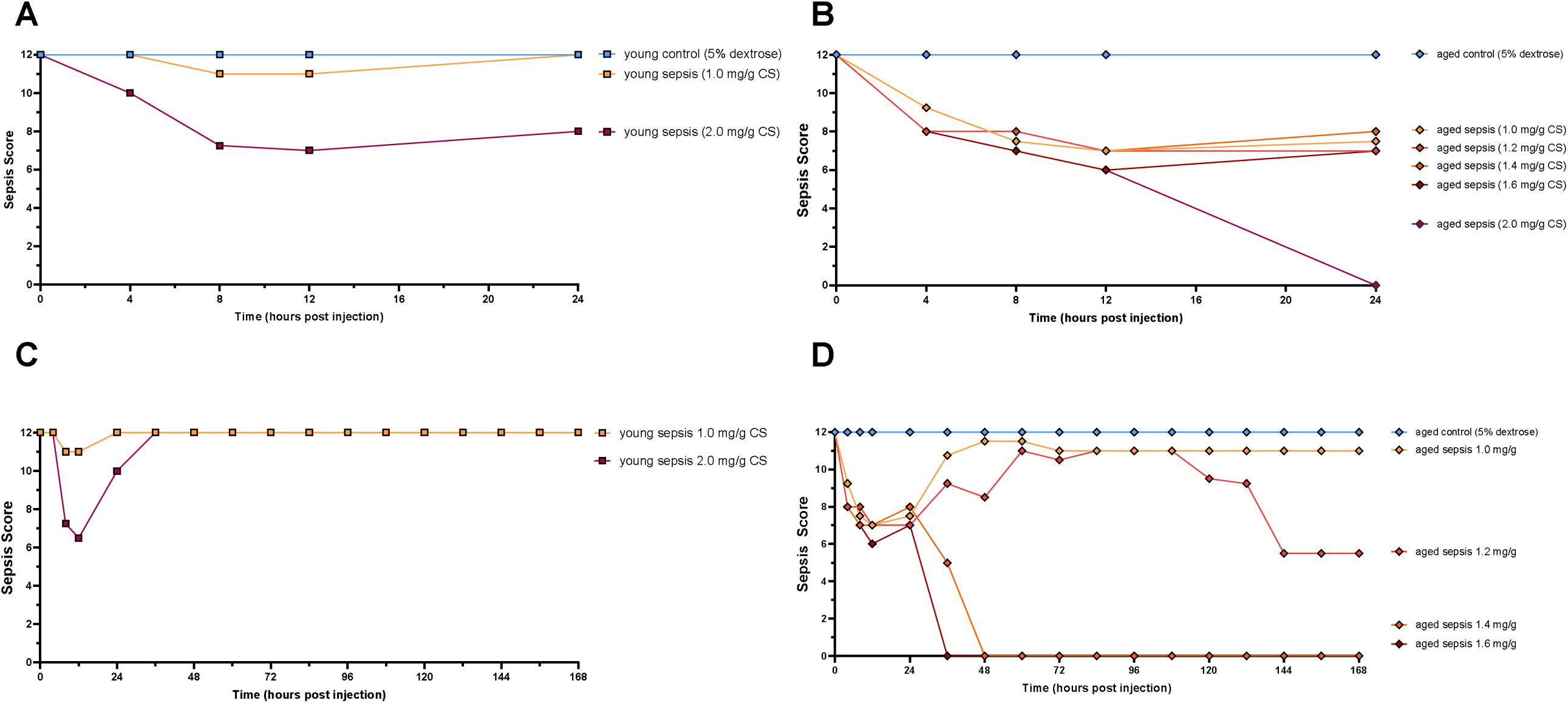
Severity of illness in young and aged mice treated with various doses of cecal slurry (CS). Both young and aged mice experienced increased illness severity with increasing doses of CS. Young mice treated with 1.0 mg/g CS only showed a brief pattern of mild sickness that resolved by 24 hours, and a dose of 2.0 mg/g CS was necessary to manifest a more severe form of illness (A). However, the 2.0 mg/g dose for 24 hours was fatal in aged mice (B), and various doses between 1.0 mg/g and 2.0 mg/g had to be tested to ensure appreciable illness without excessive fatality. Similarly, in the recovery model, young mice treated with 1.0 mg/g CS experienced mild illness and recovered by 24 hours while those treated with 2.0 mg/g CS experienced more severe illness and did not fully recover until 36-hours after injection (C). However, again, doses higher than 1.0 mg/g were fatal to aged mice over the course of 7 days after injection of 1.0 mg/g CS (D). Data were presented as median only (A, C) or mean only (B, D). N = 4-9.

**Supplemental Figure 2.**
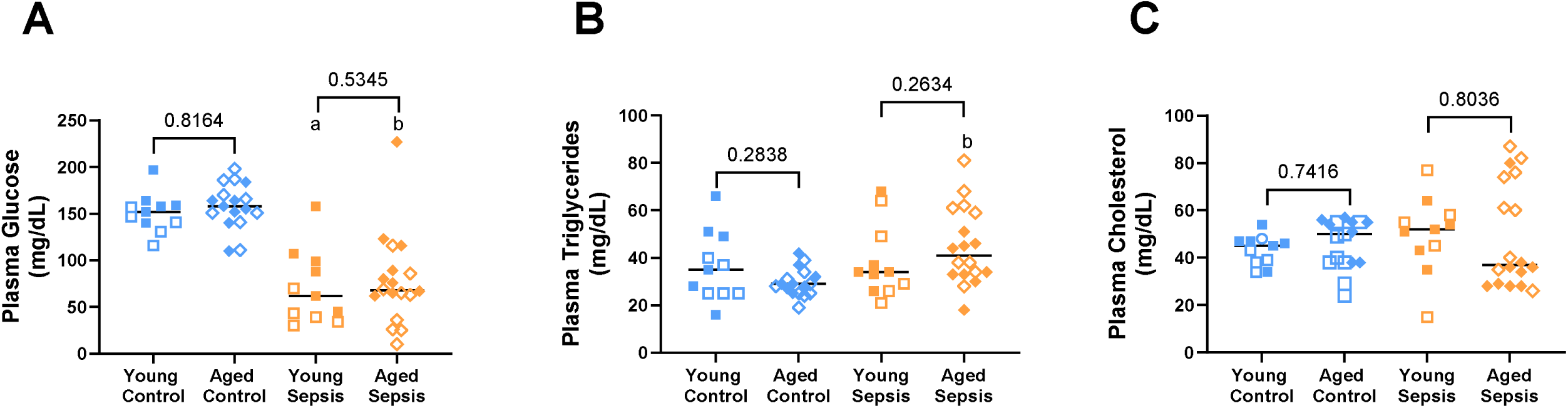
Blood glucose, blood triglycerides, and blood cholesterol associated with 24-hour septic insult [1.6 mg/g cecal slurry (CS)] in young and aged mice. Both young and aged septic mice showed decreased plasma glucose compared to their respective control mice (A). Only aged septic mice showed an increase in plasma triglyceride concentrations compared to control mice (B). Plasma cholesterol concentrations were not affected by sepsis or age (C). Data were presented as individual data points each representing an individual male (solid) or female (open) animal (A-C). Horizontal line indicates combined median for male and female animals. N = 11-18. [Statistical analysis: Two-way ANOVA + Uncorrected Fisher’s LSD test on log-transformed data (A-C).] ap<0.05 vs Young Control; bp<0.05 vs Aged Control; *p<0.05 Young Sepsis vs Aged Sepsis. Control = 5% dextrose, Sepsis = 1.6 mg/g CS.

**Supplemental Figure 3.**
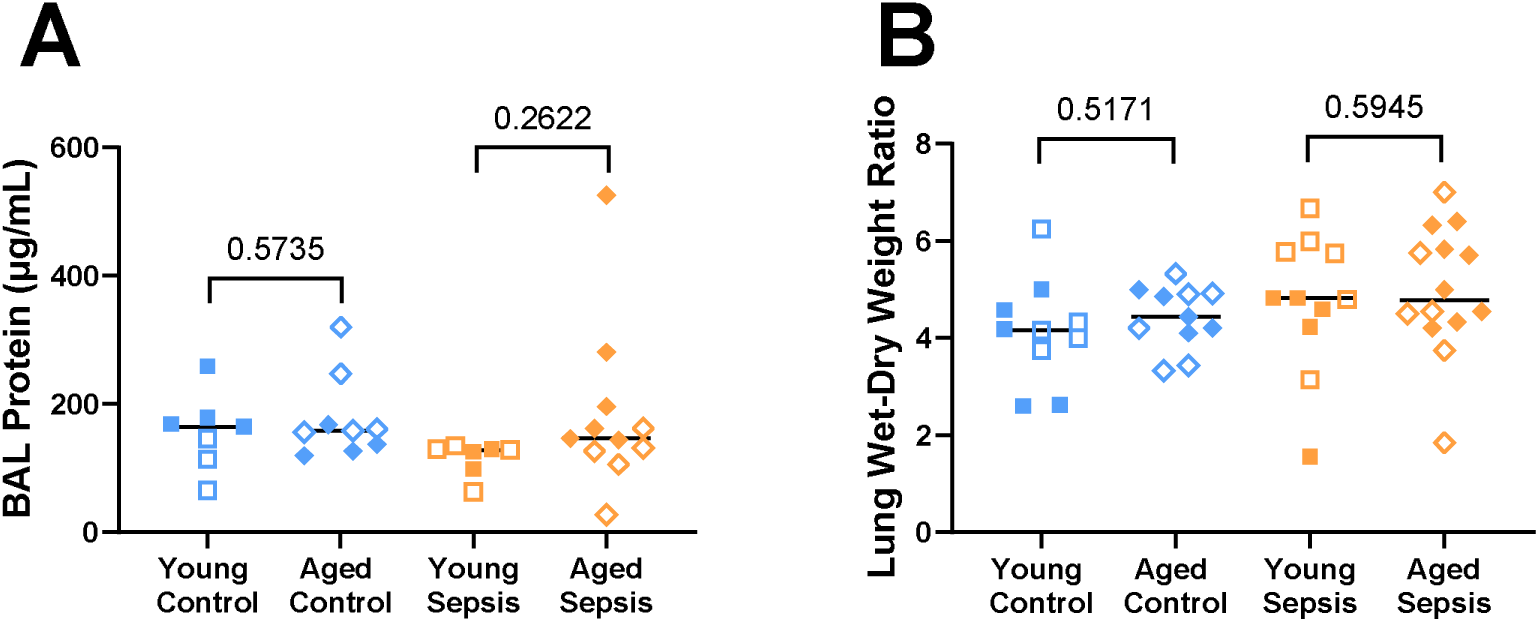
Alveolar-capillary barrier permeability associated with 24-hour septic insult [cecal slurry (CS; 1.6mg/g)] in young and aged mice. Sepsis and age had no effect on BAL protein content (A) or lung wet-to-dry ratios (B). Each point represents an individual male (solid circle) or female (open circle) animal (A-B). Horizontal line indicates combined median for male and female animals. N = 7-11. [Statistical analysis: Two-way ANOVA + Uncorrected Fisher’s LSD test on log-transformed data (A-B).] ap<0.05 vs Young Control; bp<0.05 vs Aged Control. BAL = bronchoalveolar lavage, Control = 5% dextrose, Sepsis = 1.6 mg/g CS.

**Supplemental Figure 4.**
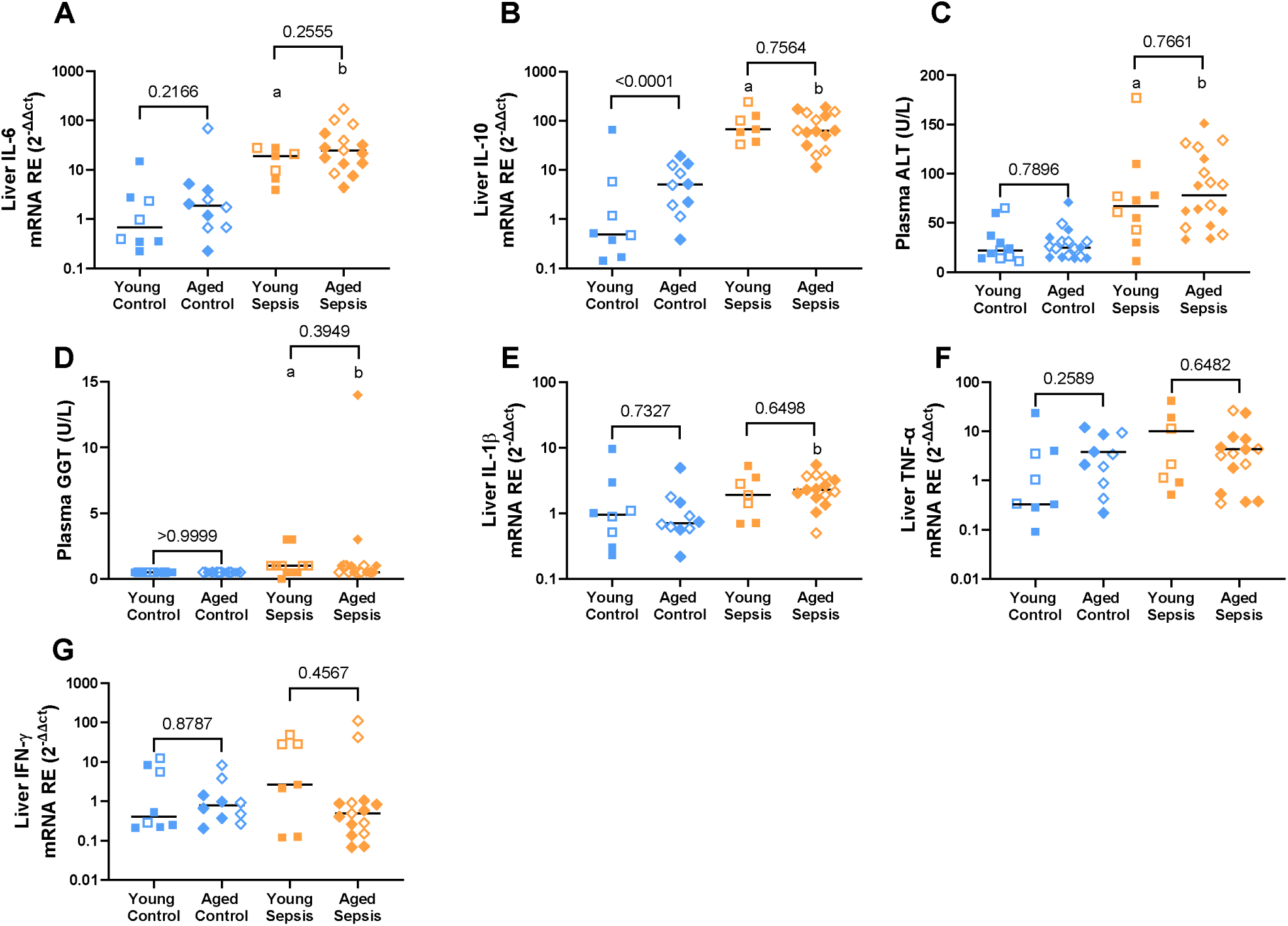
Liver inflammation and injury associated with 24-hour septic insult [cecal slurry (CS; 1.6mg/g)] in young and aged mice. Young and aged septic mice had higher liver tissue mRNA expression for IL-6 (A) and IL-10 (B), as well as plasma ALT (C) and GGT (D) compared to their respective controls. Only aged septic mice had higher liver tissue mRNA expression of IL-1β (E) compared to control mice. No differences were observed between treatment or age group for liver tissue mRNA expression of TNF-α (F) or IFN-γ (G). Each point represents an individual male (solid) or female (open) animal (A-G). Horizontal line indicates combined median for male and female animals. N = 7-18. [Statistical analysis: Two-way ANOVA + Uncorrected Fisher’s LSD test on log-transformed data (A-G)]. ap<0.05 vs Young Control; bp<0.05 vs Aged Control. Control = 5% dextrose, Sepsis = 1.6 mg/g CS, IL-6 = interleukin-6, IL-10 = interleukin-10, ALT = alanine aminotransferase, GGT = gamma-glutamyl transferase, IL-1β = interleukin-1 beta, TNF-α = tumor necrosis factor alpha, IFN-γ = interferon gamma.

**Supplemental Figure 5.**
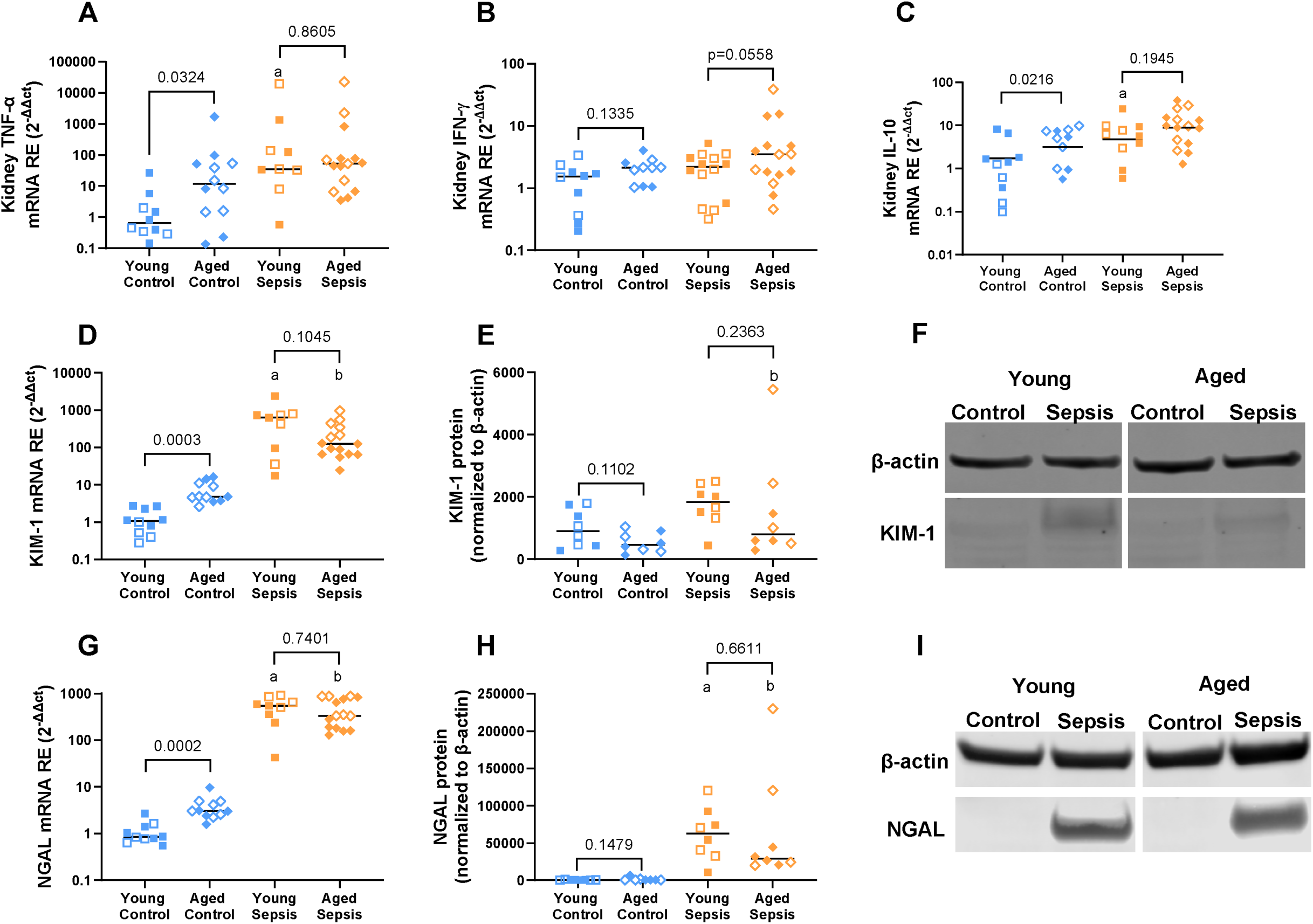
Kidney inflammation and injury associated with 24-hour septic insult [cecal slurry (CS; 1.6mg/g)] in young and aged mice. Septic aged mice had elevated kidney tissue KIM-1 mRNA (D), KIM-1 protein (E), NGAL mRNA (G), and NGAL protein (H), as well as numerically higher TNF-α (p=0.0637; A), IFN-g (p=0.1816; B), and IL-10 (p=0.0566; C) compared to aged control mice. Similar increases were observed in young septic mice compared to young control mice, except for expression of kidney tissue KIM-1 protein. At baseline, aged control mice also showed increased KIM-1 and NGAL mRNA expression in kidney tissue compared to young control mice. Compared to young septic mice, aged septic mice had numerically higher IFN-g and IL-10. However, no differences were observed for kidney tissue mRNA expression of TNF-a, or any markers of kidney injury between young and aged septic mice. Representative images of KIM-1 (F) and NGAL (I) Western blots. Each point represents an individual male (solid) or female (open) animal (A-E, G-H). Horizontal line indicates combined median for male and female animals. N = 8-15. [Statistical analysis: Two-way ANOVA + Uncorrected Fisher’s LSD test on log-transformed data (A-E, G-H).] ap<0.05 vs Young Control; bp<0.05 vs Aged Control. Control = 5% dextrose, Sepsis = 1.6 mg/g cecal slurry, TNF-α = tumor necrosis factor alpha, IFN-γ = interferon gamma, IL-10 = interleukin-10, KIM-1 = kidney injury molecule-1, NGAL = neutrophil gelatinase associated lipocalin.

**Supplemental Figure 6.**
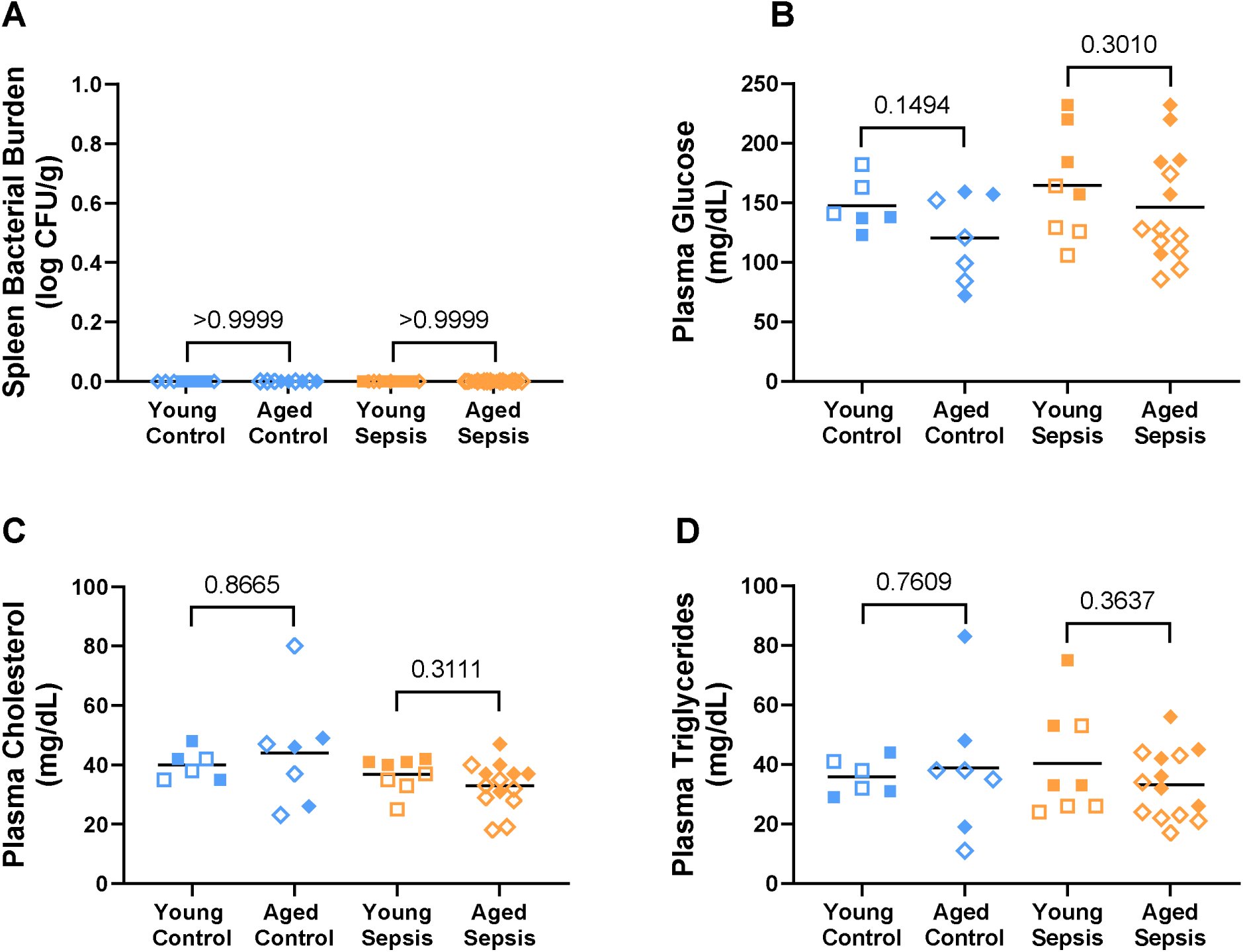
Bacterial burden, blood glucose, blood triglycerides, and blood cholesterol associated with 8-day recovery from septic insult [1.0 mg/g cecal slurry (CS)] in young and aged mice. No differences were observed between septic or control mice in either age group for spleen bacterial counts (A), plasma glucose concentrations (B), plasma cholesterol concentrations (C), or plasma triglyceride concentrations (D). However, plasma glucose was decreased in aged septic and aged control mice compared to their young counterparts. Data were presented as individual data points each representing male (solid) or female (open) animal (A-D). Horizontal line indicates combined median for male and female animals. N = 6-19. [Statistical analysis: Kruskal-Wallis test + Dunn’s multiple comparisons test (A), Two-way ANOVA + Uncorrected Fisher’s LSD test (B-D).] ap<0.05 vs Young Control; bp<0.05 vs Aged Control. Control = 5% dextrose, Sepsis = 1.0 mg/g cecal slurry.

**Supplemental Figure 7.**
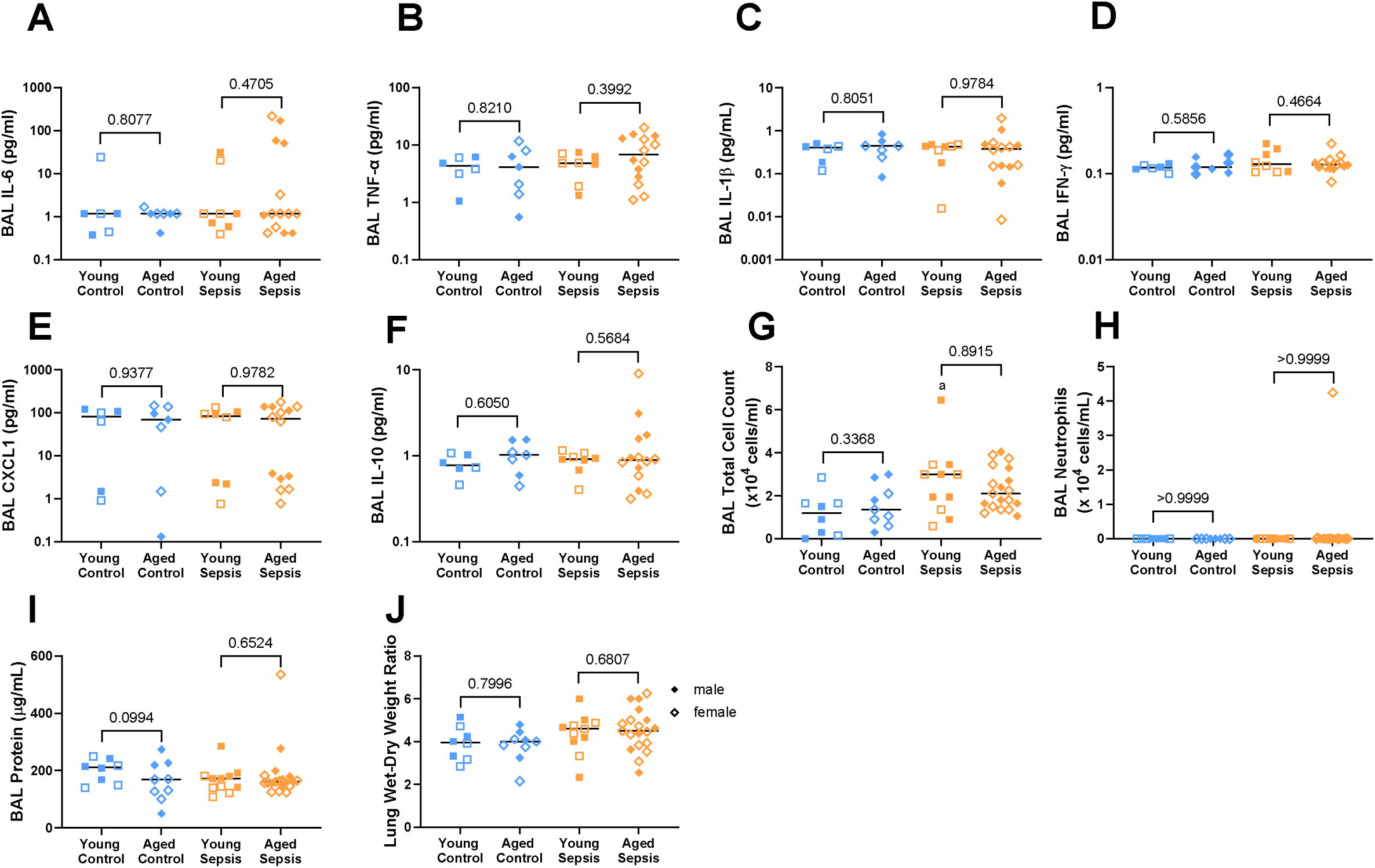
Lung inflammation and alveolar-capillary barrier permeability associated with 8-day recovery from septic insult [1.0 mg/g cecal slurry (CS)] in young and aged mice. Septic mice (both young and aged) did not differ compared to control mice for BAL concentrations of IL-6 (A), TNF-α (B), IL-1β (C), IFN-γ (D), CXCL1 (E), or IL-10 (F), BAL neutrophils (H), BAL protein concentrations (I), and lung wet-to-dry weight ratios (J). However, young septic mice had higher BAL inflammatory cell counts compared to young control mice, while aged septic mice had numerically higher BAL inflammatory cell counts compared to aged control mice (p=0.0592; G). No differences were observed between young and aged septic mice across any of the markers measured. Each data point represents an individual male (solid circle) or female (open circle) animal (A-J). Horizontal line indicates the combined median of male and female animals. N = 6-19. [Statistical analysis: Two-way ANOVA + Uncorrected Fisher’s LSD test on log-transformed data (A-G, I-J), Kruskal-Wallis test + Dunn’s multiple comparisons test (H).] ap<0.05 vs Young Control; bp<0.05 vs Aged Control. Control = 5% dextrose, Sepsis = 1.0 mg/g cecal slurry, BAL = bronchoalveolar lavage, IL-6 = interleukin-6, TNF-α = tumor necrosis factor alpha, IL-1β = interleukin-1 beta, IFN-γ = interferon gamma, CXCL1 = C-X-C motif chemokine ligand 1, IL-10 = interleukin-10.

**Supplemental Figure 8.**
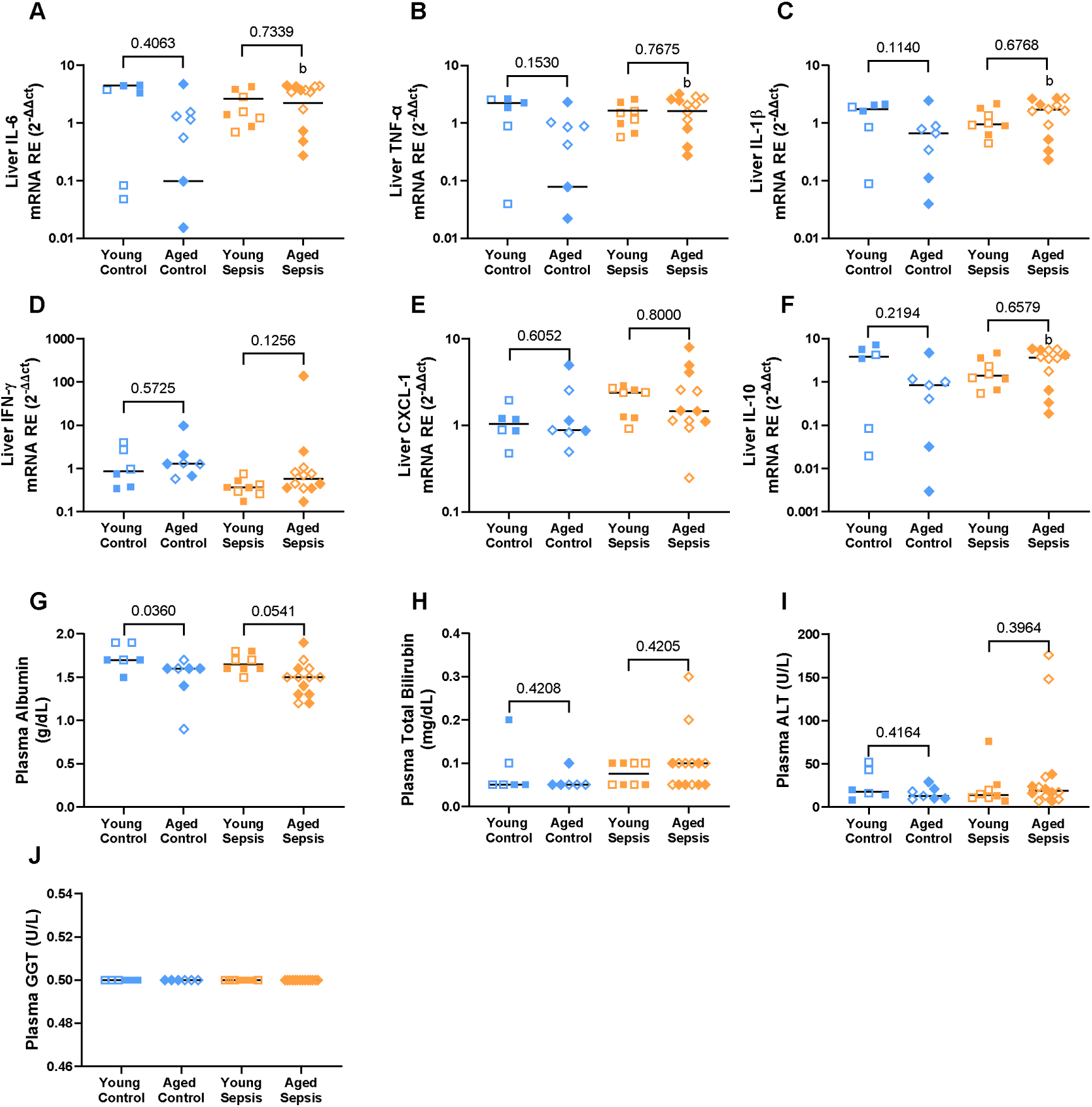
Liver inflammation, dysfunction and injury associated with 8-day recovery from septic insult [1.0 mg/g cecal slurry (CS)] in young and aged mice. Liver tissue mRNA expression for IL-6 (A), TNF-α (B), IL-1β (C), and IL-10 (F) were higher in aged septic mice compared to aged control, but not different between young septic and young control mice. Compared to their respective controls, young and aged septic mice did not differ in liver tissue mRNA expression of IFN-γ (D) or CXCL1 (E). However, aged septic mice had numerically higher IFN-γ compared to young septic mice. Septic mice (both young and aged) did not differ from control mice in plasma concentrations of albumin (G), total bilirubin (H), ALT (I), or GGT (J). However, aged control had lower plasma albumin compared to young control mice, while septic aged mice had numerically lower plasma albumin compared to young septic mice. Each data point represents an individual male (solid) or female (open) animal (A-D). Horizontal line indicates the combined median of male and female animals. N = 6-14. [Statistical analysis: Two-way ANOVA + Uncorrected Fisher’s LSD test on log-transformed data (A-J)]. Control = 5% dextrose, Sepsis = 1.0 mg/g cecal slurry, IL-6 = interleukin-6, TNF-α = tumor necrosis factor alpha, IL-1β = interleukin-1 beta, IFN-γ = interferon gamma, CXCL1 = C-X-C motif chemokine ligand 1, IL-10 = interleukin-10, ALT = alanine aminotransferase, GGT = gamma-glutamyl transferase.

**Supplemental Figure 9.**
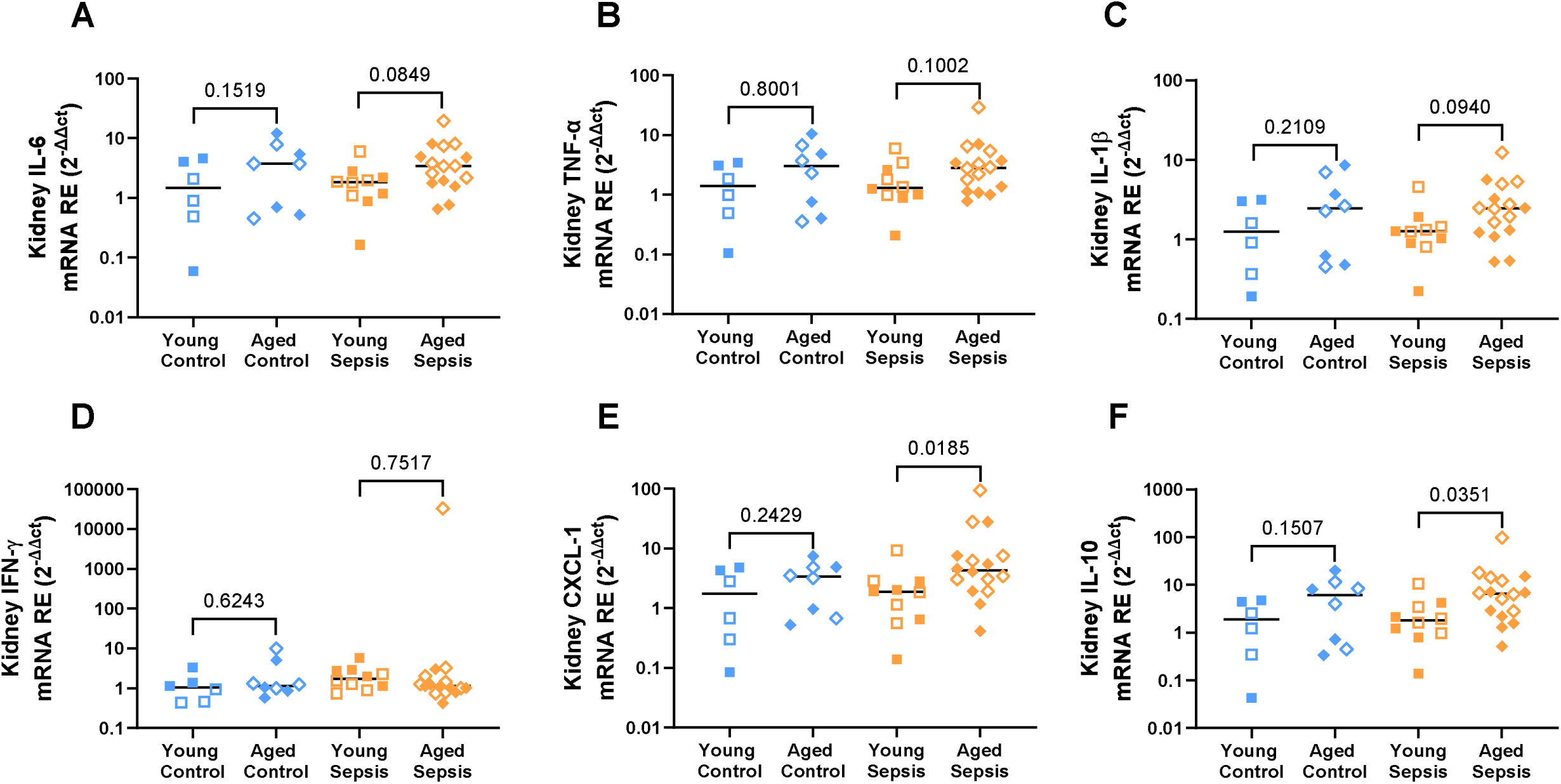
Kidney inflammation associated with 8-day recovery from septic insult [cecal slurry (CS; 1.0mg/g)] in young and aged mice. Kidney tissue mRNA expression for CXCL1 (E) was numerically higher in aged septic mice compared to aged control, but not different between young septic and young control mice. Compared to their respective controls, young and aged septic mice did not differ in kidney tissue mRNA expression of IL-10 (F), IL-6 (A), IL-1β (C), TNF-a (B), or IFN-γ (D). However, aged septic mice had higher kidney tissue mRNA expression of CXCL1 and IL-10, as well as numerically higher IL-6, IL-1β, and TNF-α compared to young septic mice. Each point represents an individual male (solid circle) or female (open circle) animal (A-F). Horizontal line indicates combined median for male and female animals. N = 6-16. [Statistical analysis: Two-way ANOVA + Uncorrected Fisher’s LSD test on log-transformed data (A-F)]. ap<0.05 vs Young Control; bp<0.05 vs Aged Control. Control = 5% dextrose, Sepsis = 1.6 mg/g CS, IL-6 = interleukin-6, TNF-α = tumor necrosis factor alpha, IL-1β = interleukin-1 beta, IFN-γ = interferon gamma, CXCL1 = C-X-C motif chemokine ligand 1, IL-10 = interleukin-10.

